# Loss of Exocytosis Protein DOC2B is an Early Event in Type 1 Diabetes Development

**DOI:** 10.64898/2025.12.28.696610

**Authors:** Diana Esparza, Eunjin Oh, Jinhee Hwang, Diti Chatterjee Bhowmick, Erika M. McCown, Jeannette Hacker-Stratton, Fouad Kandeel, Helena Reijonen, William Hagopian, Tijana Jovanovic-Talisman, Debbie C. Thurmond

**Affiliations:** Department of Molecular and Cellular Endocrinology, Arthur Riggs Diabetes & Metabolism Research Institute, Beckman Research Institute at City of Hope; Duarte, United States; Department of Translational Research and Cellular Therapeutics, Arthur Riggs Diabetes & Metabolism Research Institute, Beckman Research Institute at City of Hope; Duarte, United States; Department of Immunology & Theranostics, Arthur Riggs Diabetes & Metabolism Research Institute, Beckman Research Institute at City of Hope; Duarte, United States; Department of Medicine, University of Washington; Seattle, United States; Department of Pediatrics, Indiana University School of Medicine; Indianapolis, United States; Department of Cancer Biology and Molecular Medicine, Beckman Research Institute at City of Hope; Duarte, United States

## Abstract

**Abstract:** Type 1 diabetes (T1D) affects millions worldwide, yet few non-invasive biomarkers detect immune-mediated β-cell dysfunction during the presymptomatic phase, a critical window for therapeutic intervention. Previously, we identified reduced double C2-like domain containing beta protein (DOC2B) levels in circulating platelets as a marker of reduced β-cell function in early-onset T1D cohorts and nonobese diabetic (NOD) mice. Here, we assessed whether plasma DOC2B could serve as a sensitive early biomarker of T1D progression risk in the autoantibody-positive pediatric cohort (progressors vs non-progressors) from the longitudinal Diabetes Evaluation in Washington (DEW-IT) study; T1D patients and non-diabetic cohorts from the DEW-IT study served as controls. At pre-onset, progressors showed a decline in DOC2B that preceded measurable changes in random C-peptide and HbA1c levels, while non-progressors maintained stable levels. These observations were further supported by our analysis in prediabetic NOD mice. Comparisons of plasma levels pre- and post-clinical islet transplantation in long-standing T1D patients highlights its potential utility as a reporter of β-cell functional mass. Together, these findings suggest that DOC2B decline may precede C-peptide decline in early presymptomatic T1D progression. This work could have significant future implications for clinical trial stratification and assessing response outcomes to disease-modifying or cell replacement therapies.

## INTRODUCTION

Type 1 diabetes (T1D) is an autoimmune disease characterized by destruction of insulin-producing pancreatic β-cells, resulting in insulin deficiency (*1*). Furthermore, interventions that preserve or restore β-cells are likely to be most effective when implemented during early presymptomatic stages when insulin production is highest (*2–5*). Current predictors (e.g. genetic profiles, islet autoantibodies, and metabolic phenotypes) do not accurately capture functional β-cell decline preceding T1D onset (*6–8*), thereby limiting the therapeutic window. The development of noninvasive biomarkers of direct β-cell distress is therefore imperative for improving T1D risk prediction, clinical trial enrollment, and monitoring outcomes of disease-modifying therapies and islet transplantation.

C-peptide, secreted in a one-to-one molar ratio with insulin, is the most widely used clinical marker of β-cell activity (*9*). However, its utility is largely confined to Stage 2 T1D, when individuals with ≥ 2 islet autoantibodies exhibit subtle dysglycemia and declining C-peptide, a reflection of advanced β-cell dysfunction that fails to capture the subtle yet dynamic nature of early β-cell decline (*10, 11*). Several circulating biomarkers are under development to improve early detection of β-cell health decline (*12–15*). Elevated proinsulin:C-peptide ratios [an indicators of β-cell endoplasmic reticulum (ER) stress] and cell-free demethylated insulin DNA (an indicator of β-cell apoptosis) have been validated in autoantibody-positive individuals prior to clinical onset (*4, 16–19*). However, early T1D pathogenesis includes impaired stimulus-coupled vesicle trafficking for insulin secretion, a process regulated by soluble N-ethylmaleimide-sensitive factor attachment protein receptor (SNARE) complex proteins. Current markers do not capture these mechanisms (*20–22*). In this context, the double C2-like containing beta (DOC2B) protein may offer added value as a novel biomarker reflecting this disruption.

DOC2B is a SNARE accessory protein, involved in membrane dynamics and secretory granule trafficking, essential for biphasic glucose-stimulated insulin secretion and β-cell survival under cytokine-stress (*23–29*). In clinical onset cohorts, DOC2B levels decline in blood-derived platelets and in insulin-positive islet cells and remain low after glucose normalization through insulin therapy (*30*), indicating its independence from exogenous insulin-mediated glycemic control. In T1D models, nonobese diabetic (NOD) mice, DOC2B loss occurred before glycemic changes (*30*), supporting its potential as an early biomarker of β-cell dysfunction. However, because platelets are scarce in repositories and may reflect acute rather than chronic physiological states (*31*), we investigated whether plasma DOC2B could serve as a more accessible and clinically relevant biomarker.

Here, we hypothesized that reduced plasma DOC2B levels reflect early pre-onset, β-cell dysfunction and may serve as sensitive indicators of T1D risk and progression. To test this, we assessed DOC2B levels in plasma from the following: 1) pediatric autoantibody-positive individuals in the Diabetes Evaluation in Washington (DEW-IT) study who did or did not progress to T1D, compared to individuals with T1D (0-12.6 year disease duration) or without diabetes; 2) adult patients with long-standing, severe T1D (10-52 year disease duration) before and after clinical islet transplantation to assess dynamic changes; 3) young presymptomatic NOD mice to compare DOC2B and C-peptide loss during early disease. We found that plasma DOC2B levels predicted and distinguished presymptomatic autoantibody-positive progressors from non-progressors, reflected changes in functional β-cell mass, and declined before clinical onset in the NOD mouse model. These results position plasma DOC2B as a versatile, non-invasive tool for monitoring residual and grafted islet function, including stem cell-derived or conventional transplants, and for stratifying patients into clinical trials and guiding therapeutic interventions.

## RESULTS

### Subhead 1: Plasma double c2-like domain beta protein levels correlate with islet β-cell function

Our previous work involving two patients with long-standing T1D undergoing clinical islet transplantation, demonstrated that DOC2B protein levels in blood-derived platelets correlate with functional β-cell mass (*30*). Given the limited availability of platelets in repositories, we first used our in house DOC2B antibody, recently validated in (*32*), to examine DOC2B protein levels in both plasma and platelets from three adults with long-standing T1D (10-52 year disease duration) undergoing clinical islet transplantation as part of clinical trials [NCT01909245] and [NCT03746769] at City of Hope (COH), previously described in (*30, 33*). Samples were evaluated at Day 0 (pre-) and Day 30 (post-infusion). Samples from individuals without diabetes (ND) were included (**Tables 1-3**).

**Table 1.**
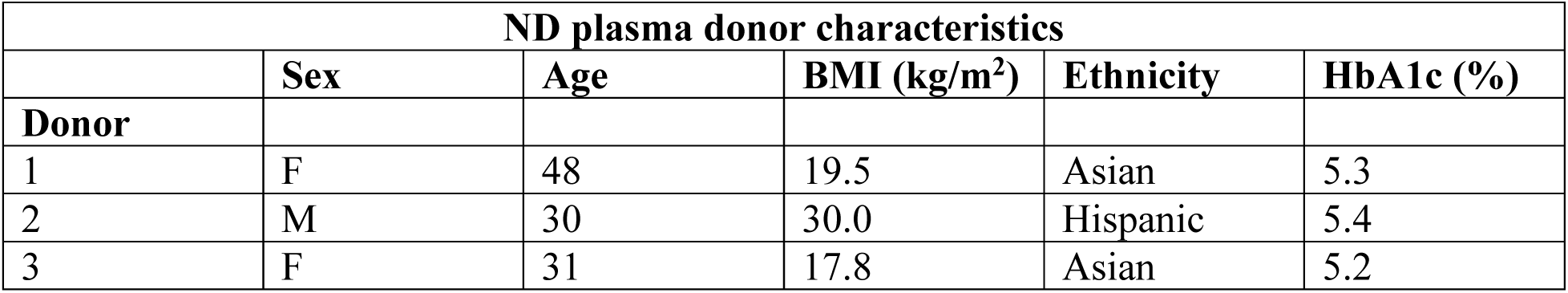
Characteristics of plasma donors without diabetes (ND) included in plasma versus platelet immunoblot analyses. .

**Table 2.** Baseline adult islet transplant recipients and islet graft characteristics and outcomes. The 1 mg intravenous (IV) glucagon stimulation test (GST) is widely used to evaluate residual β-cell insulin secretory capacity in individuals with T1D, typically under (mild) hyperglycemic conditions (*57*). This response is mediated primarily through the glucagon receptor (GLP1R) (*58*). NA, not assessed.

| Islet Transplant Recipients |  |  |  |  |  |
| --- | --- | --- | --- | --- | --- |
| Characteristics | COH-027 | COH-028 | COH-030 | COH-031 | COH-033 |
| Protocol No. | 12446 | 12446 | 18156 | 18156 | 18156 |
| <u>Recipient Baseline Characteristics</u> |  |  |  |  |  |
| Gender | F | M | F | F | M |
| Age at transplant (y) | 43 | 52 | 43 | 58 | 55 |
| Weight (kg) | 76.5 | 92.0 | 72.2 | 75.8 | 91.0 |
| BMI (kg/m<sup>2</sup>) | 28.9 | 29.8 | 28.2 | 28.2 | 26.3 |
| Duration of diabetes (years) | 33 | 34 | 27 | 52 | 33 |
| HbA1c (%) | 5.5 | 8.5 | 7.2 | 7.3 | 6.6 |
| Insulin intake (units/day) | 28 | 52 | 29 | 36 | 48 |
| Fasting/Glucagon-Stimulated C-peptide (ng/ml) | 0.03/0.02 | <0.02/<0.02 | <0.02/<0.02 | <0.02/<0.02 | <0.1/<0.1 |
| Autoantibodies | GAD65-neg, IA-2-pos, mIAA-pos, ZnT8-neg | GAD65-pos, IA-2-neg, mIAA-pos, ZnT8-neg | GAD65-neg, IA-2-neg, mIAA-pos, ZnT8-neg | GAD65-neg, IA-2-neg, mIAA-pos, ZnT8-neg | GAD65-pos, IA-2-pos, mIAA-pos, ZnT8-neg |
| Calculated PRA (%) | 22 | 0 | 0 | 0 | 0 |
| <u>Donor/Islet Graft Characteristics</u> |  |  |  |  |  |
| Islet Dose (IEQ) |  |  |  |  |  |
| Total IEQ | 240,133 | 482,755 | 370,650 | 456,683 | 344,650 |
| IEQ/kg | 3,139 | 5,247 | 5,134 | 6,025 | 3,787 |
| Islet Purity (%) | 50 | 68 | 90 | 65 | 60 |
| Packed Cell Volume (ml) | 1.9 | 2.8 | 1.2 | 2.1 | 2.0 |
| Islet Viability (%) | 91.0 | 94.0 | 95.0 | 95.0 | 84.9 |
| HbA1c (%) | 5.0 | 5.4 | 5.1 | 5.5 | 5.5 |
| BMI (kg/m<sup>2</sup>) | 36.0 | 35.9 | 39.5 | 24.0 | 28.9 |
| Age (years) | 40 | 54 | 18 | 42 | 19 |
| <u>Transplant Outcomes</u> |  |  |  |  |  |
| Insulin intake (units/day) | D30:12.5<br>D75: 15 | D30: 27.1<br>D75:19.1 | D30: 0<br>D75: NA | D30:0<br>D75: NA | D30: 0<br>D75: 0 |
|  | D183:17 | D183:12.5 | D183: 0 | D183: 0 | D183: 5 |
| **Fasting/Glucagon-stimulated C-peptide (ng/ml)** | D30:<br>2.78/3.56<br>D75:<br>0.84/1.42<br>D183:<br>0.77/1.10 | D30:<br>1.71/3.39<br>D75:<br>1.23/2.40<br>D183:<br>1.60/2.81 | D30:<br>1.85/3.4<br>D75: NA<br>D183: NA | D30:<br>2.49/6.54<br>D75: NA<br>D183: NA | D30: NA<br>D75: NA<br>D183: NA |
| **HbA1c (%)** | D75: 5.3<br>D183: 5.1 | D75: 6.0<br>D183: 6.1 | D75: 5.1<br>D183: 5.1 | D75: 5.3<br>D183: NA | D75: 6.0<br>D183: 6.5 |
| **Suspected islet graft rejection / Peak cPRA (%)** | Yes / 97 | No / 0 | No / 0 | No / 0 | No / 0 |

**Table 3.** Extended donor data set for clinical islet transplantation study. See Table 2 for details.

| **Islet Transplant Recipients** |  |  |  |  |  |
| --- | --- | --- | --- | --- | --- |
| Characteristics | **COH-034** | **COH-035** | **COH-036** | **COH-037** | **COH-038** |
| Protocol No. | **18156** | **18156** | **18156** | **18156** | **18156** |
| **<u>Recipient Baseline Characteristics</u>** |  |  |  |  |  |
| **Gender** | M | M | F | F | F |
| **Age at transplant (y)** | 55 | 37 | 31 | 37 | 52 |
| **Weight (kg)** | 76.2 | 67.7 | 61.7 | 88.7 | 72.9 |
| **BMI (kg/m<sup>2</sup>)** | 26.7 | 21.4 | 21.9 | 29.0 | 31.1 |
| **Duration of diabetes (years)** | 33 | 21 | 10 | 19 | 45 |
| **HbA1c (%)** | 7.3 | 5.8 | 6.8 | 7.4 | 6.2 |
| **Insulin intake (units/day)** | 72 | 27*<br>*low carb diet < 50g/d | 30 | 40 | 26 |
| **Fasting/Glucagon-Stimulated C-peptide (ng/ml)** | <0.1/<0.1 | <0.1/<0.1 | <0.1/<0.1 | 0.1/<0.1 | <0.1/0.1 |
| **Autoantibodies** | GAD65pos, IA-2-pos, mIAA-pos, ZnT8-neg | GAD65neg, IA-2-neg, mIAA-pos, ZnT8-neg | GAD65-neg, IA-2-neg, mIAA-pos, ZnT8-neg | GAD65-pos, IA-2-neg, mIAA-pos, ZnT8-neg | GAD65-pos, IA-2-neg, mIAA-neg, ZnT8-neg |
| **Calculated PRA (%)** | 0 | 0-56* | 0-62* | 0 | 0 |
| **<u>Donor/Islet Graft Characteristics</u>** |  |  |  |  |  |
| **Islet Dose (IEQ)** |  |  |  |  |  |
| **Total IEQ** | 362,567 | 270,292 | 363,726 | 392,925 | 291,742 |
| **IEQ/kg** | 4,758 | 3,992 | 5,895 | 4,430 | 4,002 |
| **Islet Purity (%)** | 50 | 45 | 85 (Bag 1)<br>35 (Bag 2) | 65 | 65 |
| **Packed Cell Volume (ml)** | 2.3 | 2.0 | 2.1 (Bag 1)<br>3.0 (Bag 2) | 2.2 | 2.1 |
| **Islet Viability (%)** | 96.0 | 97.5 | 94.1 (Bag 1)<br>95.6 (Bag 2) | 91.8 | 95.6 |
| **HbA1c (%)** | 5.2 | 5.3 | 5.2 | 5.1 | 5.0 |
| **BMI (kg/m<sup>2</sup>)** | 35.5 | 31.1 | 32.9 | 27.8 | 22.7 |
| **Age (years)** | 18 | 33 | 34 | 52 | 51 |
| **<u>Transplant Outcomes</u>** |  |  |  |  |  |
| **Insulin intake**<br>**(units/day)** | D30: 7.5<br>D75:8.5<br>D183: NA | D30: 16<br>D75: 13<br>D183: 22* | D30: 7<br>D75: 23<br>D183: NA | D30: 8<br>D75: NA<br>D183: 0 | D30: 0<br>D75: NA<br>D183:0 |
| **Fasting/Glucagon-**<br>**stimulated C-**<br>**peptide (ng/ml)** | D30: NA<br>D75: NA<br>D183: NA | D30: NA<br>D75: NA<br>D183: NA | D30: NA<br>D75: NA<br>D183: NA | D30: NA<br>D75: NA<br>D183: NA | D30: NA<br>D75: NA<br>D183: NA |
| **HbA1c (%)** | D75:6.2<br>D183: NA | D75: 5.5<br>D183: 6.1 | Day 75: 6.0<br>D183: NA | D75: NA<br>D183: 5.9 | D75: 5.3<br>D183: 5.9 |
| **Suspected islet**<br>**graft rejection /**<br>**cPRA (%)** | No / 0 | Yes / 99<br>GAD65-pos | Yes / 49 | No / 0 | Yes / 93 |

Immunoblot analysis validated our method to fractionate plasma from blood-derived platelets as noted by the enrichment of transferrin in the plasma fractions (**Fig. 1A-B**). Quantification revealed that independent of health status, plasma (compared to platelets) contained significantly higher levels of DOC2B per microgram of total protein (**Fig. 1A-C**). In accordance with our previous work (*30*), compared to samples from patients with long-standing T1D prior to clinical islet transplantation (Day 0), both plasma and platelets from ND donors contained higher levels of DOC2B protein (**Fig. 1A-C**). In patients with long-standing T1D, compared to Day 0 (pre-transplants), we detected significantly elevated plasma DOC2B levels on Day 30 (post-transplant); however, these levels on Day 30 were still significantly lower compared to those observed from ND donors (**Fig. 1C**). Given the plasma DOC2B abundance and good sensitivity to patient health status, we evaluated only plasma DOC2B levels in subsequent experiments.

**Fig. 1.**
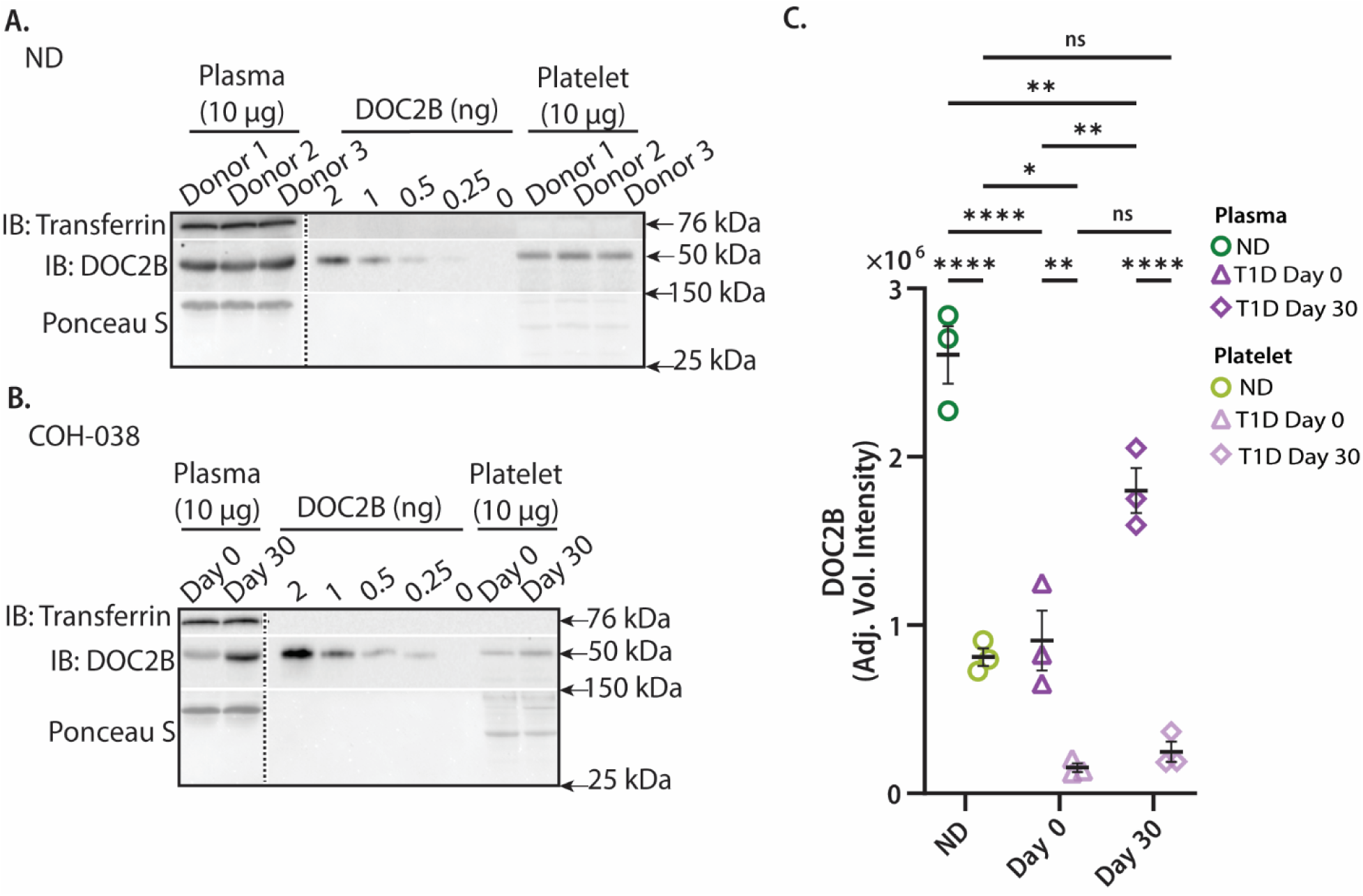
Double C2-like domain beta protein is enriched in the plasma and platelets of individuals without diabetes compared to those with T1D. Plasma and blood-derived platelets obtained from three individuals without T1D (ND) and three individuals with long-standing T1D undergoing clinical islet transplantation. Samples collected pre-islet infusion (Day 0), or 30 days after islet infusion (Day 30) were evaluated via quantitative immunoblot for DOC2B protein content. Representative immunoblots for (A) ND donors and (B) T1D participant COH-038. A dashed vertical line indicates splicing of lanes from within the same gel exposure. Ponceau S (25-150 kDa range) served as loading control. Transferrin served as a marker of plasma-enriched fractions. Recombinant DOC2B standard (2, 1, 0.5, 0.25, 0 ng) served as positive control. (C) Densitometry analysis of the DOC2B adjusted volume intensity assessed in the plasma and platelets of ND and T1D individuals (Day 0) pre- and Day 30 post-islet infusion, *n*=3 individuals per group. Data are represented as mean ± SEM.\**P* <0.05, \*\**P* <0.01, \*\*\*\**P* <0.0001, n.s. not significant, by ordinary two-way ANOVA, using Tukey’s post-hoc test.

To longitudinally assess functional changes in grafted β-cell mass *in vivo,* we immunoblotted the plasma from ten COH adult patients with long-standing T1D undergoing clinical islet transplantation (including 3 evaluated in **Fig. 1**), spanning from Day 0 up to Day 183 post-clinical islet transplantation. We consistently observed a robust elevation in plasma DOC2B levels (normalized to Ponceau S) within 30 days post-transplantation (**Fig. 2A, 2B**), further supporting our previously published results in platelets (*32*). DOC2B changes from Day 0 to Day 30 did not significantly differ in males or females (**Fig. S1A**), and did not significantly correlate with recipient age or islet graft characteristics such as total islet dose or purity; age [Pearson *r*= 0.03315, *P*= 0.9576]; total islet dose [Pearson *r*= 0.1989, *P*= 0.5818]; islet purity [Pearson *r*= -0.2497, *P*= 0.4867] (**Fig. S1B-D**).

**Fig. 2.**
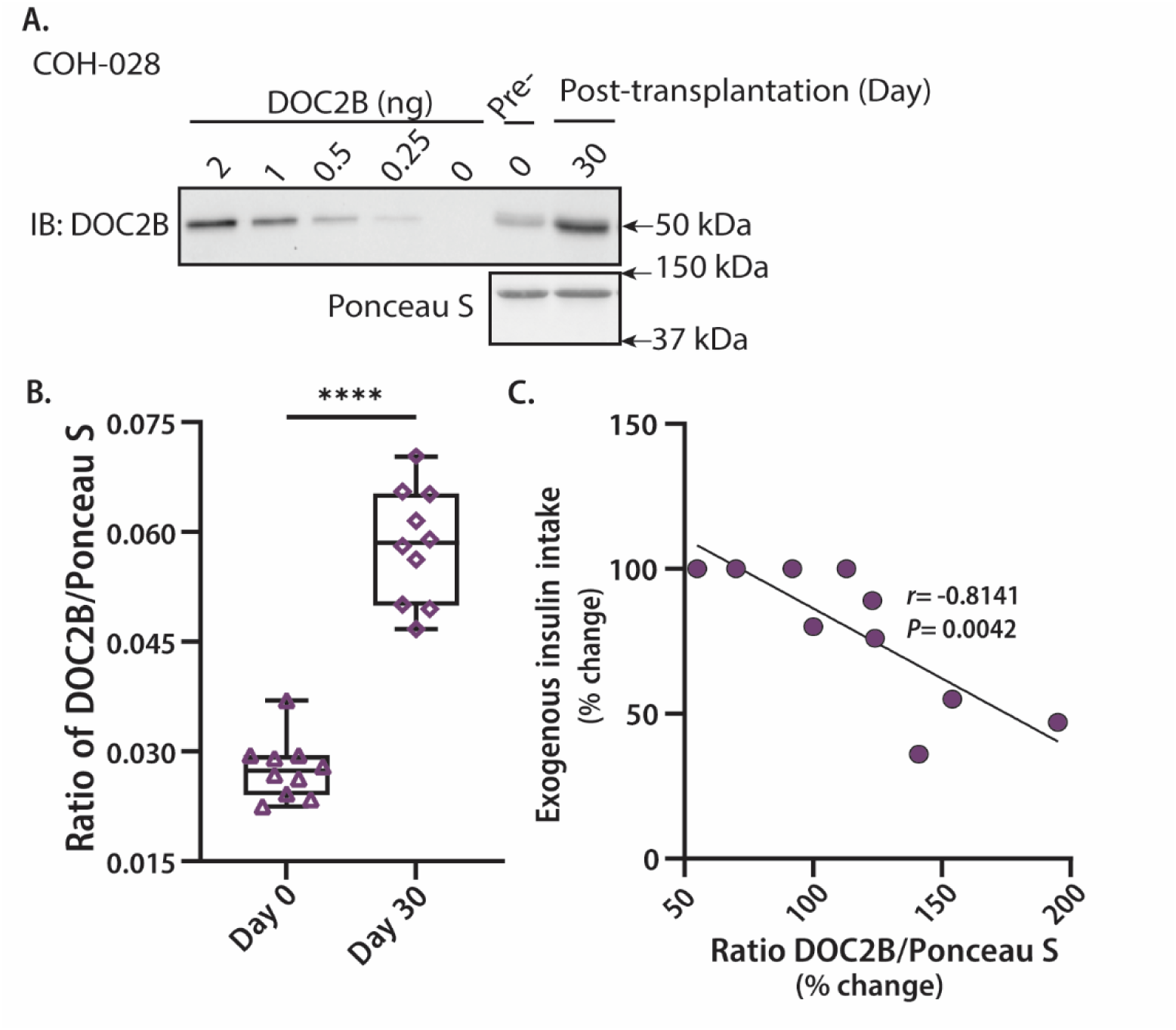
Plasma double C2-like domain beta protein as a biomarker of improved clinical outcomes in T1D. Plasma obtained from ten patients with long-standing T1D undergoing clinical islet transplantation was evaluated by quantitative immunoblotting for DOC2B protein content pre- (Day 0) islet infusion, or post-infusion: (A) Representative immunoblot of patient with long-standing T1D COH-028. Ponceau S (37-150 kDa range) served as loading control. Recombinant DOC2B standard (0-2 ng) served as positive antibody control. (B) Densitometry analysis of the ratio of DOC2B normalized to Ponceau S in the plasma of long-standing T1D patients on (Day 0; opened purple triangles), pre-, and Day 30 (opened purple diamonds) post-islet infusion, *n*=10 individuals. Data are mean ± SEM. \*\*\**P* <0.0001 by two-tailed, paired, Students *t*-test. (C) The percentage change between exogenous insulin intake and DOC2B levels (normalized to Ponceau S) on Day 30 post-versus pre-transplant (Day 0) is shown from *n*=10 individuals. There was a significant inverse correlation between the changes in exogenous insulin intake and DOC2B levels on Day 30 post-versus pre-transplant (Day 0) [Pearson *r*= -0.8141, *P*= 0042].

Elevation in plasma DOC2B on Day 30 post-transplant paralleled improvements in C-peptide and reduction in insulin requirements. C-peptide levels, measured under fasting conditions, were nearly undetectable prior to islet infusion and were restored to levels consistent within physiological range by Day 30 post-transplantation [**Tables 2-3, Fig. S2A**; data available for four participants]. Exogenous insulin therapy requirements decreased across all patients within 30 days post-transplantation compared to pre-transplant levels (**Tables 2-3, Fig. S2B**). Correlation analysis of Day 30 vs. Day 0 data revealed a statistically significant inverse relationship between insulin intake and DOC2B levels [*r*= -0.8141, *P*=0.0042] (**Fig. 2C**), an association not observed with islet graft characteristics, including total islet dose [Pearson *r*= 0.1989, *P*= 0.5818] and islet viability [Pearson *r*= 0.2311, *P*= 0.5205] (**Fig. S2C-D**).

Assessment of additional time points post-transplant revealed that relative to baseline (Day 0), plasma DOC2B levels remained elevated at Day 75 and decreased at Day 183 (**Fig. S3**; patients resuming insulin therapy after Day 30 are highlighted). Longitudinal analysis was limited to Day 183 due to protocol-related interventions: four patients experienced islet graft rejection with donor-specific antibodies, two of which were given Rituxan (COH-027 and COH-038), while COH-027 received additional immunomodulatory treatments [e.g. immunoglobulin (IVIG) and steroids, plasmapheresis; Supplementary Materials and Methods, **Tables 2-3**], and eight patients underwent immunosuppressive regimen adjustments (e.g. sirolimus reduction and tacrolimus increase) for a second gastrin-17 treatment. These interventions occurred close to the time of blood collection for subsequent plasma DOC2B evaluation, and one patient (COH-036) discontinued participation before Day 183, reducing sample availability. Given the small number of participants with complete post-rejection data and non-uniform gastrin administration (two patients did not receive it), analyses were performed across all transplant recipients without stratification by graft outcome or gastrin exposure.

### Subhead 2: Reduction in circulating double c2-like domain-containing beta protein occurs earlier than decline of C-peptide levels and hyperglycemia in NOD Mice

In our previous study, we observed a significant reduction of DOC2B protein levels in the islet cell lysates from female non-obese diabetic (NOD) mice as early as 7 weeks of age (*30*), a time point close to when pancreatic islet β-cells undergo senescence and experience functional decline (*34*). Therefore, we investigated whether DOC2B deficiency is reflected in both plasma and in insulin-positive islet cells, in parallel with degree of insulitis (monocyte infiltration) typically observed in NOD mice at this age.

We first examined plasma DOC2B protein levels in 7-week-old female NOD mice and age-, sex-, major histocompatibility complex-matched non-obese diabetes-resistant (NOR) mice relative to C-peptide, the biomarker of standard of care. Plasma DOC2B levels were decreased by at least 20% in NOD mice compared with NOR (p< 0.05) (**Fig. 3A-B**). At this age, NOR and NOD mice showed similar glucose levels (165 ± 12 mg/dl versus 140 ± 3 mg/dl, respectively) (**Fig. 3C**), consistent with these model phenotypes at 7 weeks of age. C-peptide levels showed no significant differences between NOR and NOD mice (**Fig. 3D**).

**Fig. 3.**
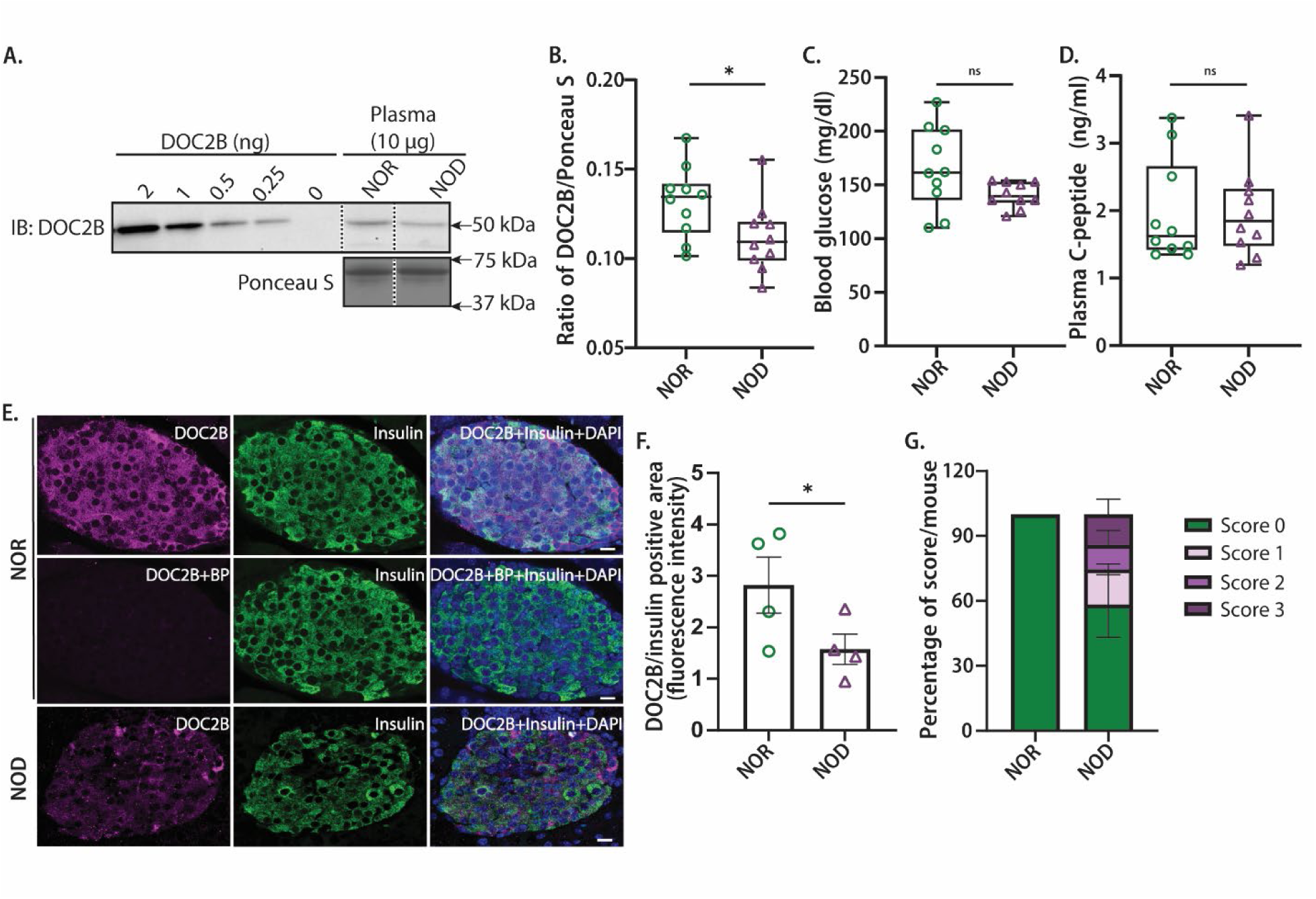
Reduced plasma double C2-like domain beta protein levels precede hallmarks of T1D in pre-onset NOD mice. Plasma from 7-week-old female nonobese diabetes (NOD) and age-, gender, and MHC-matched nonobese diabetes resistant (NOR) mice was evaluated by quantitative immunoblotting for DOC2B protein content. (A) A **r**epresentative immunoblot is shown from *n*=10 mice per group. Dashed vertical lines indicate splicing of lanes from within the same gel exposure. Ponceau S served as loading control (37-75 kDa range). (B) Densitometry analysis of the ratio of plasma DOC2B levels (normalized to Ponceau S) in the 7-week-old female NOR versus NOD. (C) Random fasting blood glucose levels were measured in NOR versus NOD mice at 7 weeks of age. (D) Plasma C-peptide levels were assessed using an enzyme-linked immunosorbent assay (ELISA). Data are represented as mean ± SEM; *n*=10 mice per group; \**P*<0.05 and n.s.-not significant using one-tailed, unpaired, Student’s *t*-test for (B), and two-tailed, unpaired, Student’s *t*-test for [C, D]. (E) DOC2B (magenta), without or with blocking peptide (BP), and insulin (green) immunofluorescence staining; (F) bar graph quantification of DOC2B fluorescence intensity in insulin-positive cells in mouse pancreata, representative of *n*=4 mice per group. Bar= 10 µm. Merge image contains DOC2B (magenta), insulin (green) and DAPI (blue), nuclear staining. Data represent the mean ± SEM. \**P*<0.05 by one-tailed, unpaired, Student’s *t*-test. (G) Quantification of the insulitis scores from hematoxylin-insulin-counterstained pancreatic tissue sections of 7-week-old female NOD and NOR mice, as described in Supplemental Materials and Methods. Data are represented as average mean ± SEM, *n*=4 mice per group.

To determine whether DOC2B deficiency is present in insulin-positive β-cells, we evaluated DOC2B protein levels in 7-week-old NOD mice and NOR mouse islets via immunofluorescence imaging (**Fig. 3E**). The specificity of our antibody was validated by the loss of DOC2B signal in the NOR islets in the presence of a blocking peptide targeting the antibody’s epitopes (**Fig. 3E**). According to measurements of relative immunofluorescent intensities, DOC2B levels were significantly reduced in insulin-positive cells of the NOD mouse islets compared to the NOR controls (**Fig. 3F**), suggesting a potential link between DOC2B deficiency and ongoing β-cell stress/dysfunction in the NOD model at this age.

To determine the level of immune cell infiltration into the pancreatic islets of Langerhans, hematoxylin and insulin counterstaining were performed on paraffin-embedded pancreata sections from 7-week-old NOD versus NOR female mice via immunohistochemistry (**Fig. S4** and **Fig. 3G**). NOR mice had largely undetectable levels of insulitis (nearly 100%=stage 0), while the insulitis scoring in NOD mice at this early age was heterogeneous: >50% of the islets exhibited no cell infiltration (score=0), while <20% showed peri-insulitis-invasive insulitis (scores between 1-3 range), respectively (**Fig. S4**). No significant differences were observed in the percentage of islets displaying peri-infiltrates, infiltrating cells extending the peri-islet, or invasive insulitis between NOD and NOR mice (**Fig. 3G**). These data suggest that plasma DOC2B protein level reduction precedes C-peptide decline and onset of hyperglycemia, highlighting its potential as an early marker of β-cell dysfunction.

### Subhead 3: Double c2-like domain beta protein stratifies autoantibody-positive individuals at risk for type 1 diabetes progression

We next sought to determine whether plasma DOC2B levels are associated with pre-onset β-cell dysfunction in T1D. To this end, we evaluated DOC2B content in plasma samples collected prior to determination of disease onset from an autoantibody-positive pediatric cohort, and compared these to two separate controls: patients with T1D (0-12.6 years disease duration) and ND controls from the Diabetes Evaluation in Washington (DEW-IT) study (*35*) [**Fig. 4**].

**Fig. 4.**
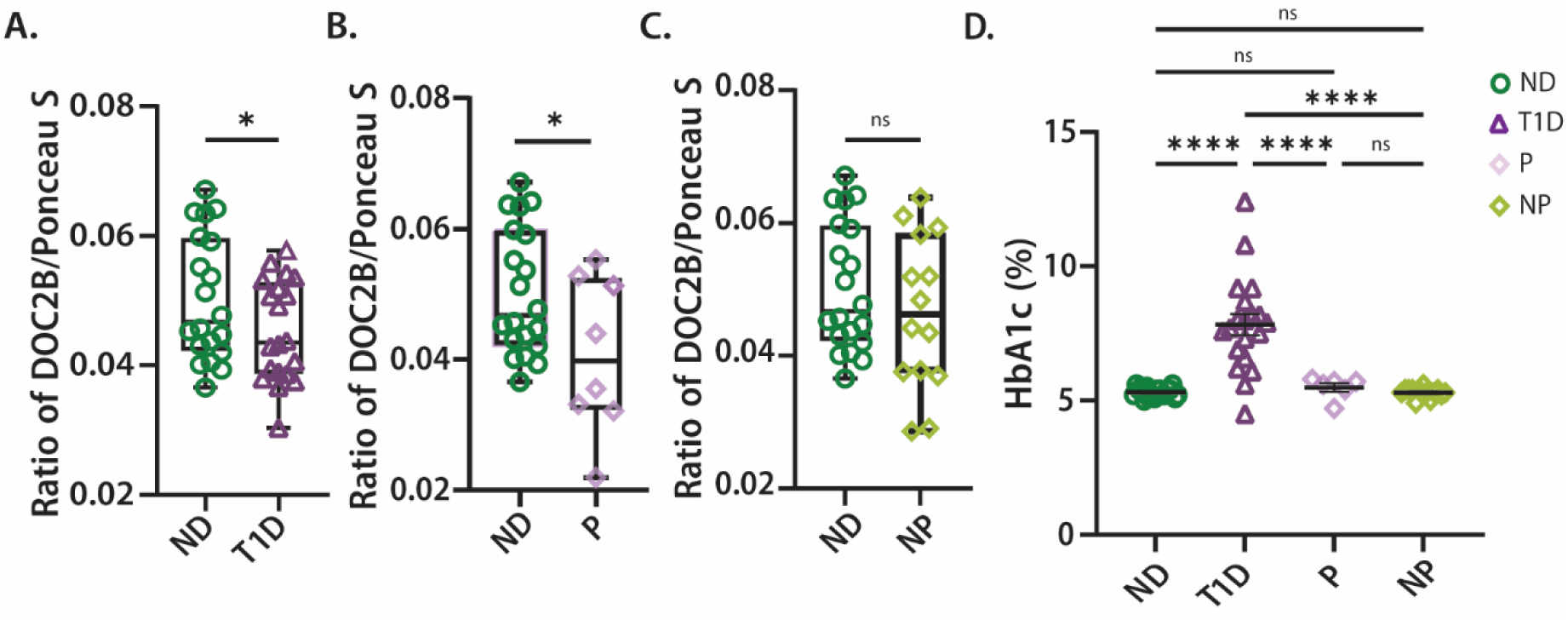
Loss of double C2-like domain beta protein serves as a biomarker for risk stratification in pre-onset type 1 diabetes. Plasma proteins from individuals with T1D and individuals without diabetes (ND), autoantibody-positive participants who progressed to T1D (P) and those who did not progress (NP) were evaluated by quantitative immunoblot for DOC2B content; *n*=62 total plasma samples evaluated. Densitometry analysis of the ratio of DOC2B normalized to Ponceau S (37-150 kDa range) for (A) ND versus T1D participants, (B) ND versus P, (C) ND versus NP. Data are represented as mean ± standard error of the mean (SEM); *\*P*<0.05 and n.s.-not significant by unpaired one-tailed Student’s *t*-test. (D) Percent HbA1c levels in ND, T1D, P, NP, *n*=62 plasma samples evaluated as independent replicates. Data are represented as mean ± SEM; *\*\*\*\*P*<0.0001 and n.s-not significant by ordinary one-way ANOVA, using Tukey’s post-hoc test.

Neither gender nor age significantly influenced DOC2B levels in non-progressors, and control patients with T1D and ND groups (**Fig. S5A-E**). Consistent with our prior detection in platelets of new-onset pediatric patients (*30*) and data in **Fig. 1**, relative to ND controls, DOC2B levels were significantly reduced in control patients with T1D (**Fig. 4A**). When compared to ND controls, DOC2B levels were significantly lower in autoantibody-positive patients who progressed to T1D diagnosis within 6-32 months (average 16 months, median 14.6 months) after sampling, but not in non-progressors to T1D across the 60-month follow-up period (**Fig. 4B-C**, **Table 4)**. Importantly, both autoantibody-positive progressors and non-progressors exhibited HbA1c levels that did not significantly differ from ND controls, whereas control patients with T1D displayed significantly elevated HbA1c levels (**Fig. 4D**). Intriguingly, there was no difference in average random C-peptide levels at the time of sampling among autoantibody-positive progressors and non-progressors, despite the significant difference in DOC2B levels, whereas control patients with T1D displayed lower levels (**Table 4**).

**Table 4.**
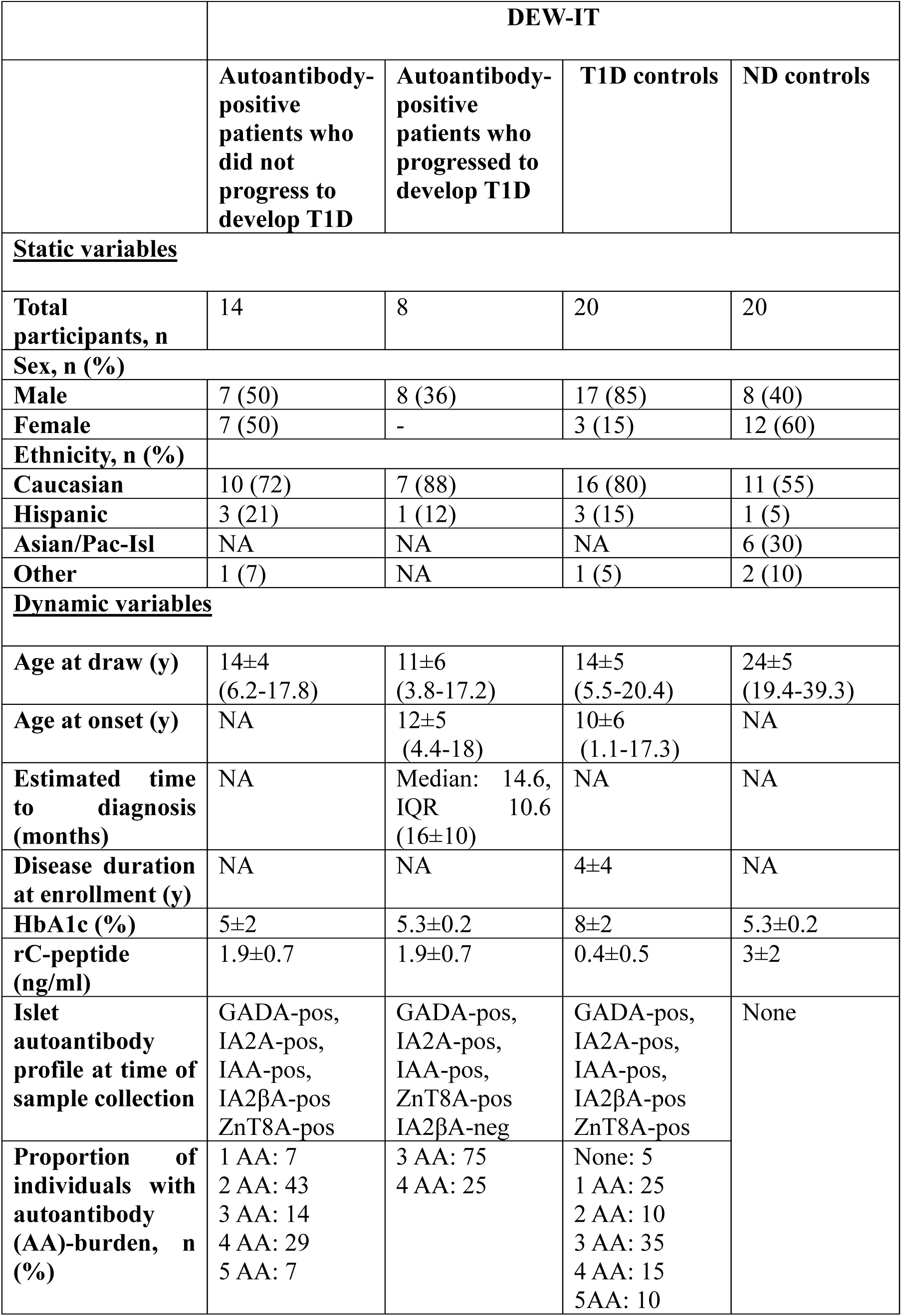
Demographic information for the Diabetes Evaluation in Washington (DEW-IT) cohort. ND= individuals without diabetes, r=random, Mean ± SD, IQR=interquartile range. Estimated time to diagnosis was determined by subtracting age at onset from age of drawn, multiplied by 12 for months in a year.

## DISCUSSION

Identifying biomarkers that reflect early, intrinsic β-cell changes during the presymptomatic phase of T1D is critical: 1) to enable timely interventions that may delay onset and reduce disease burden in at-risk individuals; and 2) to improve the design and enrollment criteria for clinical trials. Due to the silent and gradual development of T1D, few molecular signatures are detectable in blood prior to disease onset that inform on β-cell functional changes. Autoantibodies help identify individuals at risk at Stage 1, but their presence neither guarantees progression to Stage 3, disease onset, nor provide direct insight into ongoing β-cell functional decline (*36*). Current staging biomarkers, such as HbA1c, glycemia and C-peptide, are informative for Stage 2, when dysglycemia emerges, but have limited utility for Stage 1 (*37*). C-peptide, in particular, requires stimulated or carefully timed sampling (e.g. typically during a mixed-meal tolerance test) and can be challenging to interpret, and gold-standard methods like glycemic clamps are too complex for routine use (*37–39*).

Elevated circulating proinsulin:C-peptide ratios are emerging indicators of declining functional β-cell mass in individuals with multiple autoantibody-positivity and those who develop T1D, compared to ND controls (*18, 40*). Recent findings suggest that a high proinsulin:C-peptide ratio in Stage 2 T1D (pre-onset) may help predict progression to Stage 3 (T1D onset) by a median of 35.4 months in advance, and can also serve as readout for disease-modifying therapies such as Teplizumab, which delayed onset to a median of 59.7 months in responders (*4*). Our findings suggest that plasma DOC2B levels emerge as an additional promising biomarker of functional β-cell mass, and may carry promise for even earlier detection and intervention.

In the clinical COH cohort of adults with long-standing T1D undergoing islet transplantation, baseline (Day 0) plasma DOC2B levels were significantly lower (mean 0.028, SEM 0.001) compared to levels observed in pediatric T1D patients from the DEW-IT study (mean 0.045, SEM 0.002) [**Fig. S6**]. Notably, DOC2B levels in the COH cohort rose to levels above those noted in DEW-IT study ND individuals (mean 0.050 SEM, 0.002) by Day 30 post-transplantation (mean 0.058, SEM 0.002). Thereafter, in four COH cohort patients, plasma DOC2B levels declined by Day 75 (mean 0.040, SEM 0.005) and approached baseline by Day 183 (mean 0.030, SEM 0.002) in six patients. Indeed, this observed increase in plasma DOC2B levels at Day 30 post-transplant coincided with elevated fasting C-peptide levels and significantly reduced insulin intake, indicating robust functionality of the islet graft. This suggests that early post-transplant β-cells may rely on DOC2B-regulated exocytic pathways to meet insulin demand. However, the subsequent decline in DOC2B by Day 75 and return to near-pre-transplant levels by Day 183 may reflect a normalization of cellular activity as glycemia improves post-transplant. Intriguingly, a concurrent decline in plasma DOC2B occurred in patients who resumed exogenous insulin therapy, a trend evident even in those with limited sampling on these timepoints. In one case (COH-033), loss of plasma DOC2B preceded the need for return to insulin therapy by 108 days. Given that there are few noninvasive biomarkers of β-cell function that are truly useful when crucially required, such as Stages 1 and 2 T1D, or monitoring transplanted islets, our findings herein support the concept of plasma DOC2B fulfilling these unmet needs as a biomarker of islet β-cell function.

Further analysis in islet clinical transplantation recipients is needed to confirm this observation and to explore the predictive potential of DOC2B. For example, in the case of allogeneic transplant rejection, the immune response is directed to the foreign tissue of the donor, not β-cells harboring specific defects. Therefore, plasma DOC2B measurements may give insight into the predictive capacity of DOC2B to detect changes in β-cell functionality before C-peptide loss in other scenarios, such as autoimmune reactivation β-cell stress/exhaustion. It remains unclear whether the additional immune interventions these patients received to halt islet rejection (e.g., plasmapheresis) could have impacted DOC2B presentation in plasma.

Our examination of the pre-onset T1D model (NOD mice) at the time of emergent insulitis and β-cell stress provide the first evidence that plasma DOC2B levels decline before measurable C-peptide loss and dysregulated glycemic control. This aligns with prior work demonstrating that early alterations in pancreatic islet architecture, signaling, and inflammatory responses occur in pre-T1D, while changes in C-peptide are not yet detectable despite ongoing β-cell stress in NOD mice (*41, 42*). In line with this, our study demonstrates, for the first time, the predictive capacity of plasma DOC2B in distinguishing presymptomatic autoantibody-positive progressors from non-progressors. In this study, progressors exhibited lower plasma DOC2B levels (mean 0.041, SEM 0.004) as early as 6-32 months (average 16 months, median 14.6 months) after sampling, prior to clinical onset of T1D (**Tables 4-5**), 20.8 months median lead time over that of when increased proinsulin:C-peptide ratios are reported to occur, while maintaining HbA1c and random C-peptide levels comparable to ND controls. These findings suggest that DOC2B loss precedes decline in C-peptide and overt hyperglycemia, and may help identify individuals who require immediate interventions versus those that do not, even in the absence of hyperglycemia. These findings highlight DOC2B as a superior and sensitive biomarker for detecting early changes in β-cell function. Indeed, DOC2B’s consistent reduced circulating abundance in T1D individuals, across multiple cohorts, including individuals with new-onset, early-onset T1D and individuals with long-standing T1D in (*30*), further characterized here, and now pre-onset stages, supports its relevance for disease stratification.

While our study provides novel insights into DOC2B as a potential early prognostic biomarker for T1D, some limitations warrant consideration. First, although plasma DOC2B levels correlated with β-cell dysfunction, the precise source and release mechanisms remain unclear. Plasma is a complex biofluid and DOC2B may exist in soluble form or associated with extracellular vesicles (EVs), as we previously demonstrated in β-cell derived EVs (*32*). Second, the DEW-IT study was limited by the number of autoantibody-positive progressors and non-progressors compared to both onset T1D and ND controls, reducing statistical power. Future studies of individuals with genetic risk or first-degree relatives sampled prior to autoimmunity onset will be valuable toward determining the capacity of DOC2B to reflect intrinsic β-cell changes before immune attack. If validated, DOC2B could complement existing biomarkers to identify high-risk individuals earlier and guide interventions to preserve β-cell function. Finally, interpretation of Day 183 post-transplant data is confounded by adjustments in immunosuppressive therapy (e.g. sirolimus reduction and tacrolimus increase) administered alongside a second dose of gastrin to promote β-cell regeneration. These changes may alter immune balance and β-cell stress, while hypoxia associated with transplantation could further contribute to DOC2B decline toward basal levels (*43, 44*).

Our findings highlight that plasma DOC2B analysis may add value to current early T1D assessments, particularly to determine which autoantibody-positive individuals may be at higher risk to progress to disease, possibly with longer lead time before measurable β-cell loss. This strategy could widen the window of opportunity for timely intervention to delay/prevent T1D clinical progression. Future studies aimed at validating DOC2B as a biomarker and exploring its role in β-cell function could pave the way for improved early detection and targeted therapeutic interventions in T1D.

## MATERIALS AND METHODS

### Subhead 1: Study design

This study investigated the capacity of circulating DOC2B levels in reflecting early β-cell functional distress in the pre-onset stages of T1D compared to C-peptide, in clinically viable liquid biopsies. Using plasma from patient cohorts at varying stages of T1D, including those with autoantibody-positivity, sampled before disease progression, as well as those with long-standing T1D undergoing clinical islet transplantation. We analyzed plasma and histological findings in murine pancreatic islets to validate experimental findings.

The number of animals used to test the hypothesis was determined on the basis of variance from previous findings (*30, 45*). Data collection was terminated when the planned number of animals or pancreatic islet studies and replicates had been performed, based on comparisons in the “Statistical analysis” section. All data, including outliers, were analyzed. Genotype information was concealed from investigators during data acquisition and analysis of in vivo experiments (blood glucose, C-peptide, insulitis scoring) and immunoblot of plasma samples. Upon quantification of the data for each sample, the clinical team reidentified samples to permit grouping of data into T1D, ND, autoantibody-positive progress and non-progressors for participants from the DEW-IT study, and shared clinical outcomes of long-standing T1D islet transplantation patients for statistical comparisons. The clinical outcomes for COH-027 and COH-028 Day 0, Day 30 and Day 75, were known per (*30*).

### Subhead 2: Animals

Animals were maintained under protocols approved by the City of Hope (Duarte, CA USA) Institutional Animal Care and Use Committee (IACUC) Protocol No. 23042 and following the National Research Council Guidelines for the Care and Use of Laboratory Animals. Six-week-old female non-obese diabetic (NOD; NOD/ShiLtJ; RRID: IMSR_JAX:001976) and major histocompatibility complex (MHC)-matched control non-obese diabetes-resistant (NOR; RRID:I MSR_JAX:002050) mice were obtained from the Jackson Laboratory (Bar Harbor, ME, USA) and were housed in a room with a 12 hour light/dark cycle. At seven weeks of age, morning blood glucose levels were measured at 0800 hour using the Alphatrak2 glucose meter from Zoetis (Parsippany, NJ, USA), and then whole blood was collected from the retro-orbital sinus of NOD and NOR mice using EDTA-coated whole blood tubes. Following plasma separation, C-peptide levels were measured using a mouse C-peptide ELISA kit purchased from ALPCO (Salem, NH, USA, cat# 80-cptms-e01), according to the manufacturer’s instruction. Plasma and platelet isolation from whole blood is described in Supplemental Materials and Methods.

### Subhead 3: Human participants

To evaluate DOC2B content in both plasma and platelet components of blood, samples from individuals without diabetes (ND); donors 1, 2, 3, were obtained under approval from the City of Hope Institutional Review Board IRB Protocol No.18318. Three participants, aged 30-48, were recruited based on HbA1c levels <5.7% (see **Table 1** for demographic data), and blood was collected using acid citrate dextrose (ACD)-coated whole blood tubes.

To evaluate DOC2B levels in both plasma and platelet components of blood of individuals with T1D from the islet transplantation study, samples were obtained from T1D islet transplantation recipients at City of Hope, as approved by the City of Hope Institutional Review Board IRB Protocol No. 12446 [NCT01909245] and IRB Protocol No. 18156 [NCT03746769]. This cohort has been previously described in (*30, 33*).Ten participants, aged 31-58 years, were recruited for human islet transplantation based on the following criteria: Adults (Age 18-68 years), T1D for at least five years, absence of β-cell function (C-peptide level of ≤ 0.2 ng/ml before and ≤0.3 ng/ml after IV administration of 1 mg of glucagon), with frequent or life-threatening hypoglycemia, hypoglycemia unawareness and/or otherwise unstable blood glucose control. Each participant received a single islet transplant as described in Supplemental Materials and Methods. Whole blood was obtained from all participants before transplantation (Day 0) and on follow-up appointments, among them Day: 30, 75, 91, 183 after islet transplantation (see **Table 2-3** for demographic data). Blood samples were initially collected in tubes coated with ACD and later in tubes containing EDTA due to supply shortages during the COVID-19 pandemic. Prior proteomics studies indicate that the core platelet proteome remains largely consistent across commonly used anticoagulants, provided samples are processed within 1-2 hours (*46*), as done in our study. Plasma and platelet isolation from whole blood is described in Supplemental Materials and Methods.

To determine DOC2B levels preceding T1D diagnosis, human plasma from participants in the Diabetes Evaluation in Washington (DEW-IT) study was obtained from the Pacific Northwest Research Institute as approved by the City of Hope Institutional Review Board IRB Protocol No.18318 not human participant research, a cohort previously described in (*35*). In this observational study, participants born in Washington State between 2000-2004, from a general population identified through newborn screening, carrying increased risk genotype were HLA-DQ-DR-DQ genotypes and first-degree family relatives of people with T1D (irrespective of HLA genotype) were eligible for further studying based on the criteria: having at least one or two high-risk HLA haplotypes (DQA1*03-DQB1*03:02 and/or DQA1*05:01-DQB1*02:01) plus no protective haplotype (DQA1*01-DQB1*05:03, DQA1*01-DQB1*06:01, DQA1*01-DQB1*06:02, DQA1*01-DQB1*06:03, DQA1*05-DQB1*0301, DQA1*03-DQB1*03:01, DQA1*02-DQB1*03:03 or DQA1*A2-DQB1*02:02). Eligible children were offered periodic surveillance for islet autoantibodies (GADA, IA-2A, IAA, ZnT8A) by mail-based or provider-based sampling via testing in a DASP/IASP participating laboratory under CLIA certification (CLIA#50D0982418) as described in (*47*). Children with no autoantibodies were followed less frequently. After ∼15 years follow-up from the confirmed initial detection of islet autoantibodies, T1D development was confirmed in a subset of participants presenting autoantibody-positivity at the time of initial detection. Samples were collected in tubes coated with EDTA and shipped to the clinical site at ambient temperature, processed within 24 hours of draw, and stored at -80 °C. Samples were thawed only once onto wet ice for aliquoting, refrozen at -80 °C and shipped overnight on dry ice for our studies. Demographic information pertaining to these individuals is included in **Table 4**. Clinical/laboratory assays are described in detail in Supplemental Materials and Methods.

### Subhead 4: Antibody-based methods

Immunoblot was performed as described below. Detailed protocols for immunofluorescence staining and immunohistochemistry, and insulitis scoring thereof, are provided in the Supplemental Materials and Methods.

### Immunoblot

Mouse or human plasma was centrifuged at 18,900 xg for 5 minutes at 4 °C to remove debris. The samples were lysed in 1% NP40 buffer at a 1:20 dilution. DC protein assay reagents purchased from Bio-Rad laboratories (Hercules, CA, USA, cat# 5000113, 5000114, 5000115) were used to quantify protein concentration of human plasma (1:20 diluted) and human platelet samples per manufacturer’s instructions. Mouse plasma protein concentrations were determined by Bradford’s dye-binding method using a BSA standard (*48*). Plasma and platelet proteins (10 µg) were resolved onto a 10% or a 15% SDS-PAGE and transferred onto 0.2 µm polyvinylidene fluoride (PVDF) membrane (BioRad laboratories, cat# 1620177) for immunoblotting.

The membranes were blocked with 50 mg/ml non-fat milk in TBST (0.1% Tween 20) for 1 hour at room temperature. After blocking, membranes were washed five times with TBST at 1-minute intervals. All primary antibodies were diluted in TBST supplemented with 10 mg/ml BSA and 0.02% sodium azide for use on membranes. Primary antibodies used for these experiments include the in-house DOC2B antibody #2, extensively validated in (*32*), and the commercially available antibody against Transferrin from Abcam (Waltham, MA, USA, cat# ab109503) to validate these as plasma samples. Primary antibodies diluted 1:2000 for anti-DOC2B and 1:30,000 for anti-Transferrin were applied onto membranes for overnight incubation at 4 °C. Following the primary antibody incubation, the membranes were washed three times at 10 minute-intervals with TBST. Membranes were subsequently incubated for 1 hour at room temperature with horseradish peroxidase-conjugated goat anti-rabbit IgG (HL) (Bio-Rad laboratories, cat# 172-1019) secondary antibody; the secondary antibody were prepared in blocking solution and diluted 1:5000 for anti-DOC2B or 1:20,000 for anti-Transferrin. Following the secondary antibody incubation, the membranes were washed three times at 10 minute-intervals with TBST. Immunoreactive bands were visualized with Amersham ECL Western Blotting Detection Reagent from GE Healthcare (Chicago, IL, USA, cat# RPN2106), Amersham ECL prime Detection Reagent (GE Healthcare, cat# RPN2232), and SuperSignal West Femto Chemiluminescent substrate (Thermo Fisher Scientific, cat# 34095) and imaged using the ChemiDoc gel documentation system (Bio-Rad laboratories). Immunoblot exposures for DOC2B analysis were normalized for protein loading in each well by Ponceau S, as done previously in (*30*). Upon quantification of the data for each sample, the clinical team re-identified samples to permit grouping into categories for the DEW-IT cohort: control T1D patients, ND controls, autoantibody-positive progressors versus non-progressors. These groupings enabled statistical comparisons and integration with clinical outcomes. Likewise, clinical outcomes for T1D participants in the islet transplantation cohort were revealed upon data quantification and re-identification, allowing for correlation with DOC2B levels. Of note, outcomes for COH-027 and COH-028 for Day 0, 30 and 75 were known for these patients due to findings reported in (*30*).

### Subhead 5: Statistical analysis

Data are expressed as the average mean ± standard error of the mean (SEM). Unpaired Student’s *t*-test was performed for two groups of samples to compare DOC2B levels in the plasma of DEW-IT T1D patients, autoantibody-positive progressors, or autoantibody-positive non-progressors with ND individuals, as well as for DOC2B levels in plasma, random blood glucose, plasma C-peptide, DOC2B/ insulin area fluorescence intensity data from 7-week-old female NOD versus NOR mice. A one-tailed test was performed, where indicated, based on a priori hypothesis predicting the direction of the effect. Specifically, prior evidence suggests that DOC2B levels would be lower in individuals with T1D compared to those without diabetes (*30*). Therefore, a one-tailed test was deemed appropriate to increase statistical power for detecting effect in the hypothesized direction, while acknowledging that effects in the opposite direction would not be considered significant under this framework.

Paired Student’s *t*-test was performed for two groups of samples to compare plasma DOC2B levels and exogenous insulin intake in long-standing T1D participant samples on Day 0, pre-, and Day 30, post-islet transplantation. An unpaired Welch’s test was performed to compare fasted C-peptide levels on Day 0, pre-, and Day 30, post-islet transplantation.

An ordinary two-way ANOVA with Tukey’s post-hoc test was performed to compare plasma DOC2B levels in males versus females from DEW-IT ND and T1D individuals, as well as to compare DOC2B levels in the plasma and platelets from ND donors or in those with long-standing T1D on Day 0, pre-, or Day 30 post-transplantation. An ordinary one-way ANOVA with Tukey’s post-hoc test was performed to compare HbA1c levels in DEW-IT ND, T1D, progressors, and non-progressors. All correlations were assessed with Pearson’s correlation test and simple linear regression to assess the influence of variables of interest in univariate analyses. GraphPad Prism 9. Significance was considered \**P*<0.05, \*\**P*<0.01, \*\*\**P*<0.001, \*\*\*\**P*<0.0001, n.s-not significant.

## Acknowledgments

We thank Drs. Chathurani S. Jayasena for manuscript editing, R. Vasavada, S. Shuck, and A. Garcia-Ocaña (City of Hope) for helpful discussions. We thank Dr. Arnab Chowdhury for helpful input in statistical analysis of samples, Dr. Victoria Seewaldt for sharing clinical samples to conduct this study. We also thank the blood/plasma donors and their families. Research reported in this publication included work performed in the Pathology Core, Light Microscopy/Digital Imaging Core, Biostatistics and Mathematical Oncology services, Biomarkers, Histology & Rodent Islet Isolation/Transplantation Service Center, AR-DMRI Laboratory & Translational Service Centers at the City of Hope.

## Author contributions

Conceptualization: DE, TJT, FK, HR, WH, DCT Methodology: DE, EO, JH, DCB, EM, JHS Investigation: DE, EO, JH, DCB, EM, JHS Visualization: DE, EO, JH, DCB, EM, JHS, Funding acquisition: DE, JH, DCT, TJT Project administration: DCT, TJT Supervision: TJT, FK, HR, WH, DCT Writing – original draft: DE, EO, JH, DCB, EM, JHS, TJT, HR, DCT Writing – review & editing: DE, EO, JH, DCB, EM, JHS, TJT, FK, HR, WH, DCT

## Data and materials availability

All data are available in the main text or the supplementary materials. Further inquiries can be directed to the corresponding authors.

## References

1. G. D. Ogle, F. Wang, A. Haynes, G. A. Gregory, T. W. King, K. Deng, D. Dabelea, S. James, A. J. Jenkins, X. Li, R. C. W. Ma, D. M. Maahs, R. A. Oram, C. Pihoker, J. Svensson, Z. Zhou, D. J. Magliano, J. Maniam, Global type 1 diabetes prevalence, incidence, and mortality estimates 2025: Results from the International diabetes Federation Atlas, 11th Edition, and the T1D Index Version 3.0. Diabetes. Res. Clin. Pract. 225, 112277 (2025).

2. E. K. Sims, B. N. Bundy, K. Stier, E. Serti, N. Lim, S. A. Long, S. M. Geyer, A. Moran, C. J. Greenbaum, C. Evans-Molina, K. C. Herold, Teplizumab improves and stabilizes beta cell function in antibody-positive high-risk individuals. Sci. Transl. Med. 13, (2021).

3. K. C. Herold, S. E. Gitelman, P. A. Gottlieb, L. A. Knecht, R. Raymond, E. L. Ramos, Teplizumab: A Disease-Modifying Therapy for Type 1 Diabetes That Preserves β-Cell Function. Diabetes Care. 46, 1848–1856 (2023).

4. E. K. Sims, S. M. Geyer, S. A. Long, K. C. Herold, High proinsulin:C-peptide ratio identifies individuals with stage 2 type 1 diabetes at high risk for progression to clinical diagnosis and responses to teplizumab treatment. Diabetologia. 66, 2283–2291 (2023).

5. W. E. Russell, B. N. Bundy, M. S. Anderson, L. A. Cooney, S. E. Gitelman, R. S. Goland, P. A. Gottlieb, C. J. Greenbaum, M. J. Haller, J. P. Krischer, I. M. Libman, P. S. Linsley, S. A. Long, S. M. Lord, D. J. Moore, W. V. Moore, A. M. Moran, A. B. Muir, P. Raskin, J. S. Skyler, J. M. Wentworth, D. K. Wherrett, D. M. Wilson, A.-G. Ziegler, K. C. Herold, T. D. T. S. Group, Abatacept for Delay of Type 1 Diabetes Progression in Stage 1 Relatives at Risk: A Randomized, Double-Masked, Controlled Trial. Diabetes Care. 46, 1005–1013 (2023).

6. C. J. Greenbaum, C. Speake, J. Krischer, J. Buckner, P. A. Gottlieb, D. A. Schatz, K. C. Herold, M. A. Atkinson, Strength in Numbers: Opportunities for Enhancing the Development of Effective Treatments for Type 1 Diabetes-The TrialNet Experience. Diabetes. 67, 1216–1225 (2018).

7. A. G. Ziegler, G. T. Nepom, Prediction and pathogenesis in type 1 diabetes. Immunity. 32, 468–478 (2010).

8. C. Evans-Molina, E. K. Sims, L. A. DiMeglio, H. M. Ismail, A. K. Steck, J. P. Palmer, J. P. Krischer, S. Geyer, P. Xu, J. M. Sosenko, β Cell dysfunction exists more than 5 years before type 1 diabetes diagnosis. JCI Insight. 3, (2018).

9. J. P. Palmer, G. A. Fleming, C. J. Greenbaum, K. C. Herold, L. D. Jansa, H. Kolb, J. M. Lachin, K. S. Polonsky, P. Pozzilli, J. S. Skyler, M. W. Steffes, C-peptide is the appropriate outcome measure for type 1 diabetes clinical trials to preserve beta-cell function: report of an ADA workshop, 21-22 October 2001. Diabetes. 53, 250–264 (2004).

10. P. Vardi, L. Crisa, R. A. Jackson, R. Dumont Herskowitz, J. I. Wolfsdorf, D. Einhorn, L. Linarelli, R. Dolinar, S. Wentworth, S. J. Brink, H. Starkman, J. S. Soeldner, G. S. Eisenbarth, Predictive value of intravenous glucose tolerance test insulin secretion less than or greater than the first percentile in islet cell antibody positive relatives of Type 1 (insulin-dependent) diabetic patients. Diabetologia. 34, 93–102 (1991).

11. H. P. Chase, D. D. Cuthbertson, L. M. Dolan, F. Kaufman, J. P. Krischer, D. A. Schatz, N. H. White, D. M. Wilson, J. Wolfsdorf, First-phase insulin release during the intravenous glucose tolerance test as a risk factor for type 1 diabetes. J. Pediatr. 138, 244–249 (2001).

12. E. M. Akirav, J. Lebastchi, E. M. Galvan, O. Henegariu, M. Akirav, V. Ablamunits, P. M. Lizardi, K. C. Herold, Detection of β cell death in diabetes using differentially methylated circulating DNA. Proc. Natl. Acad. Sci. U. S. A. 108, 19018–19023 (2011).

13. R. A. Watkins, C. Evans-Molina, J. K. Terrell, K. H. Day, L. Guindon, I. A. Restrepo, R. G. Mirmira, J. S. Blum, L. A. DiMeglio, Proinsulin and heat shock protein 90 as biomarkers of beta-cell stress in the early period after onset of type 1 diabetes. Transl. Res. 168, 96–106.e101 (2016).

14. R. S. Aguirre, A. Kulkarni, M. W. Becker, X. Lei, S. Sarkar, S. Ramanadham, E. A. Phelps, E. S. Nakayasu, E. K. Sims, R. G. Mirmira, Extracellular vesicles in β cell biology: Role of lipids in vesicle biogenesis, cargo, and intercellular signaling. Mol. Metab. 63, 101545 (2022).

15. S. C. Shuck, P. Achenbach, B. O. Roep, J. S. Termini, C. Hernandez-Castillo, C. Winkler, A. Weiss, A. G. Ziegler, Methylglyoxal products in pre-symptomatic type 1 diabetes. Front. Endocrinol. (Lausanne). 14, 1108910 (2023).

16. M. E. Røder, M. Knip, S. G. Hartling, J. Karjalainen, H. K. Akerblom, C. Binder, Disproportionately elevated proinsulin levels precede the onset of insulin-dependent diabetes mellitus in siblings with low first phase insulin responses. The Childhood Diabetes in Finland Study Group. J. Clin. Endocrinol. Metab. 79, 1570–1575 (1994).

17. I. Truyen, P. De Pauw, P. N. Jørgensen, C. Van Schravendijk, O. Ubani, K. Decochez, E. Vandemeulebroucke, I. Weets, R. Mao, D. G. Pipeleers, F. K. Gorus, Proinsulin levels and the proinsulin:c-peptide ratio complement autoantibody measurement for predicting type 1 diabetes. Diabetologia. 48, 2322–2329 (2005).

18. E. K. Sims, Z. Chaudhry, R. Watkins, F. Syed, J. Blum, F. Ouyang, S. M. Perkins, R. G. Mirmira, J. Sosenko, L. A. DiMeglio, C. Evans-Molina, Elevations in the Fasting Serum Proinsulin-to-C-Peptide Ratio Precede the Onset of Type 1 Diabetes. Diabetes Care. 39, 1519–1526 (2016).

19. K. C. Herold, S. Usmani-Brown, T. Ghazi, J. Lebastchi, C. A. Beam, M. D. Bellin, M. Ledizet, J. M. Sosenko, J. P. Krischer, J. P. Palmer, β cell death and dysfunction during type 1 diabetes development in at-risk individuals. J. Clin. Invest. 125, 1163–1173 (2015).

20. R. N. Bone, O. Oyebamiji, S. Talware, S. Selvaraj, P. Krishnan, F. Syed, H. Wu, C. Evans-Molina, A Computational Approach for Defining a Signature of β-Cell Golgi Stress in Diabetes. Diabetes. 69, 2364–2376 (2020).

21. M. K. Huber, A. E. Widener, A. E. Cuaycal, D. Smurlick, E. A. Butterworth, N. I. Lenchik, J. Chen, M. Beery, H. Hiller, E. Verney, I. Kusmartseva, M. S. Rupnik, M. Campbell-Thompson, I. C. Gerling, M. A. Atkinson, C. E. Mathews, E. A. Phelps, Beta cell dysfunction occurs independently of insulitis in type 1 diabetes pathogenesis. Cell. Rep. 44, 116174 (2025).

22. D. C. Thurmond, H. Y. Gaisano, Recent Insights into Beta-cell Exocytosis in Type 2 Diabetes. J. Mol. Biol. 432, 1310–1325 (2020).

23. M. Miyazaki, M. Emoto, N. Fukuda, M. Hatanaka, A. Taguchi, S. Miyamoto, Y. Tanizawa, DOC2b is a SNARE regulator of glucose-stimulated delayed insulin secretion. Biochem. Biophys. Res. Commun. 384, 461–465 (2009).

24. L. Ramalingam, E. Oh, S. M. Yoder, J. T. Brozinick, M. A. Kalwat, A. J. Groffen, M. Verhage, D. C. Thurmond, Doc2b is a key effector of insulin secretion and skeletal muscle insulin sensitivity. Diabetes. 61, 2424–2432 (2012).

25. J. Li, J. Cantley, J. G. Burchfield, C. C. Meoli, J. Stöckli, P. T. Whitworth, H. Pant, R. Chaudhuri, A. J. Groffen, M. Verhage, D. E. James, DOC2 isoforms play dual roles in insulin secretion and insulin-stimulated glucose uptake. Diabetologia. 57, 2173–2182 (2014).

26. L. Ramalingam, E. Oh, D. C. Thurmond, Doc2b enrichment enhances glucose homeostasis in mice via potentiation of insulin secretion and peripheral insulin sensitivity. Diabetologia. 57, 1476–1484 (2014).

27. D. Chatterjee Bhowmick, A. Aslamy, S. Bhattacharya, E. Oh, M. Ahn, D. C. Thurmond, DOC2b Enhances β-Cell Function via a Novel Tyrosine Phosphorylation-Dependent Mechanism. Diabetes. 71, 1246–1260 (2022).

28. A. Aslamy, E. Oh, E. M. Olson, J. Zhang, M. Ahn, A. S. M. Moin, R. Tunduguru, V. A. Salunkhe, R. Veluthakal, D. C. Thurmond, Doc2b Protects β-Cells Against Inflammatory Damage and Enhances Function. Diabetes. 67, 1332–1344 (2018).

29. D. C. Bhowmick, M. Ahn, S. Bhattacharya, A. Aslamy, D. C. Thurmond, DOC2b enrichment mitigates proinflammatory cytokine-induced CXCL10 expression by attenuating IKKβ and STAT-1 signaling in human islets. Metabolism. 164, 156132 (2025).

30. A. Aslamy, E. Oh, M. Ahn, A. S. M. Moin, M. Chang, M. Duncan, J. Hacker-Stratton, M. El-Shahawy, F. Kandeel, L. A. DiMeglio, D. C. Thurmond, Exocytosis Protein DOC2B as a Biomarker of Type 1 Diabetes. J. Clin. Endocrinol. Metab. 103, 1966–1976 (2018).

31. B. K. Manne, F. Denorme, E. A. Middleton, I. Portier, J. W. Rowley, C. Stubben, A. C. Petrey, N. D. Tolley, L. Guo, M. Cody, A. S. Weyrich, C. C. Yost, M. T. Rondina, R. A. Campbell, Platelet gene expression and function in patients with COVID-19. Blood. 136, 1317–1329 (2020).

32. D. Esparza, C. Lima, S. Abuelreich, I. Ghaeli, J. Hwang, E. Oh, A. Lenz, A. Gu, N. Jiang, F. Kandeel, D. C. Thurmond, T. Jovanovic-Talisman, Pancreatic β-cells package double C2-like domain beta protein into extracellular vesicles via tandem C2 domains. Front. Endocrinol. (Lausanne). 15, 1451279 (2024).

33. M. I. Husseiny, A. Kaye, E. Zebadua, F. Kandeel, K. Ferreri, Tissue-specific methylation of human insulin gene and PCR assay for monitoring beta cell death. PLoS One. 9, e94591 (2014).

34. P. J. Thompson, A. Shah, V. Ntranos, F. Van Gool, M. Atkinson, A. Bhushan, Targeted Elimination of Senescent Beta Cells Prevents Type 1 Diabetes. Cell. Metab. 29, 1045–1060.e1010 (2019).

35. V. Anand, Y. Li, B. Liu, M. Ghalwash, E. Koski, K. Ng, J. L. Dunne, J. Jönsson, C. Winkler, M. Knip, J. Toppari, J. Ilonen, M. B. Killian, B. I. Frohnert, M. Lundgren, A.-G. Ziegler, W. Hagopian, R. Veijola, M. Rewers, T. D. I. S. G. for the, Islet Autoimmunity and HLA Markers of Presymptomatic and Clinical Type 1 Diabetes: Joint Analyses of Prospective Cohort Studies in Finland, Germany, Sweden, and the U.S. Diabetes Care. 44, 2269–2276 (2021).

36. A. G. Ziegler, M. Rewers, O. Simell, T. Simell, J. Lempainen, A. Steck, C. Winkler, J. Ilonen, R. Veijola, M. Knip, E. Bonifacio, G. S. Eisenbarth, Seroconversion to multiple islet autoantibodies and risk of progression to diabetes in children. JAMA. 309, 2473–2479 (2013).

37. K. Vehik, D. Boulware, M. Killian, M. Rewers, R. McIndoe, J. Toppari, Å. Lernmark, B. Akolkar, A.-G. Ziegler, H. Rodriguez, D. A. Schatz, J. P. Krischer, W. Hagopian, T. T. S. Group, f. t. T. S. Group, T. S. Group, Rising Hemoglobin A1c in the Nondiabetic Range Predicts Progression of Type 1 Diabetes As Well As Oral Glucose Tolerance Tests. Diabetes Care. 45, 2342–2349 (2022).

38. R. A. DeFronzo, J. D. Tobin, R. Andres, Glucose clamp technique: a method for quantifying insulin secretion and resistance. Am. J. Physiol. Endocrinol. Metab. 237, E214 (1979).

39. R. N. Bergman, L. S. Phillips, C. Cobelli, Physiologic evaluation of factors controlling glucose tolerance in man: measurement of insulin sensitivity and beta-cell glucose sensitivity from the response to intravenous glucose. J. Clin. Invest. 68, 1456–1467 (1981).

40. T. M. Triolo, L. Pyle, S. Seligova, L. Yu, K. Simmons, P. Gottlieb, C. Evans-Molina, A. K. Steck, Proinsulin:C-peptide ratio trajectories over time in relatives at increased risk of progression to type 1 diabetes. J. Transl. Autoimmun. 4, 100089 (2021).

41. Y. Garciafigueroa, B. E. Phillips, C. Engman, M. Trucco, N. Giannoukakis, Neutrophil-Associated Inflammatory Changes in the Pre-Diabetic Pancreas of Early-Age NOD Mice. Front. Endocrinol. (Lausanne). 12, 565981 (2021).

42. A. F. Mathisen, A. M. Vacaru, L. Unger, E. M. Lamba, O.-A.-M. Mardare, L. M. Daian, L. Ghila, A.-M. Vacaru, S. Chera, Molecular profiling of NOD mouse islets reveals a novel regulator of insulitis onset. Sci Rep 14, 14669 (2024).

43. J. Lau, J. Henriksnäs, J. Svensson, P. O. Carlsson, Oxygenation of islets and its role in transplantation. Curr. Opin. Organ. Transplant. 14, 688–693 (2009).

44. K. K. Papas, T. M. Suszynski, C. K. Colton, Islet assessment for transplantation. Curr. Opin. Organ. Transplant. 14, 674–682 (2009).

45. E. Oh, E. M. McCown, M. Ahn, P. A. Garcia, S. Branciamore, S. Tang, D. F. Zeng, B. O. Roep, D. C. Thurmond, Syntaxin 4 Enrichment in β-Cells Prevents Conversion to Autoimmune Diabetes in Non-Obese Diabetic (NOD) Mice. Diabetes. 70, 2837–2849 (2021).

46. S. Tassi Yunga, A. J. Gower, A. R. Melrose, M. K. Fitzgerald, A. Rajendran, T. A. Lusardi, R. J. Armstrong, J. Minnier, K. R. Jordan, O. J. T. McCarty, L. L. David, P. A. Wilmarth, A. P. Reddy, J. E. Aslan, Effects of ex vivo blood anticoagulation and preanalytical processing time on the proteome content of platelets. J. Thromb. Haemost. 20, 1437–1450 (2022).

47. W. Woo, J. M. LaGasse, Z. Zhou, R. Patel, J. P. Palmer, H. Campus, W. A. Hagopian, A novel high-throughput method for accurate, rapid, and economical measurement of multiple type 1 diabetes autoantibodies. J. Immunol. Methods. 244, 91–103 (2000).

48. M. M. Bradford, A rapid and sensitive method for the quantitation of microgram quantities of protein utilizing the principle of protein-dye binding. Anal. Biochem. 72, 248–254 (1976).

