## supplementary materials for "Loss of Exocytosis Protein DOC2B is an Early Event in Type 1 Diabetes Development"

Diana Esparza *et al*.

The PDF file includes:

Materials and Methods

Fig. S1 to S6

References (*49-55)*

### MATERIALS AND METHODS

#### Subhead 1: Islet cell transplantation

For the T1D islet transplant study, human pancreata were procured from ABO-compatible, crossmatch negative cadaveric donors. The islets were isolated under cGMP conditions by the Southern California Islet Resource Center at City of Hope using a modified Ricordi method. Islets were transplanted intraportally with heparinized saline (35 U/kg recipient body weight) using a transhepatic percutaneous approach. All subjects received immunosuppressive induction with rabbit anti-thymocyte globulin, etanercept and anakinra and maintenance immunosuppression with a combination of low dose sirolimus, tacrolimus, and/or mycophenolate mofetil. Subjects COH-027, COH-035, COH-036, and COH-038 experienced suspected islet graft rejection shortly after transplant. COH-027 was also given intravenous immunoglobulin (IVIG), steroids, plasmapheresis and Rituxan. COH-038 was also given Rituxan. Subjects enrolled in IRB protocol No. 18156 also received investigational treatment with gastrin-17 analogue, under evaluation for improving islet transplant outcomes at City of Hope. These patients were given underwent sirolimus reduction and tacrolimus increase prior to administration of the second dose of gastrin-17.

#### Subhead 2: Clinical/laboratory assays

For the T1D islet transplant study, plasma C-peptide levels were measured with the C-Peptide II Assay Tosoh Bioscience (San Francisco, CA, USA; detection range 0.02 to 30 ng/ml) in the Northwest Lipid Metabolism and Diabetes Laboratory (Seattle, WA, USA). To confirm T1D diagnosis and absence of β-cell function before islet transplantation, subjects were confirmed to have negligible C-peptide (fasting C-peptide ≤ 0.02 ng/ml and 6-minute glucose-stimulated C-peptide ≤ 0.3 ng/ml). The levels of islet autoantibodies, including GAD-65 (GADA), IA-2A, micro-assays for insulin autoantibodies (IAA) and zinc transporter 8 (ZnT8A), were determined by radio-binding assays by the Autoantibody/HLA Service Center at the Barbara Davis Center for Diabetes (Aurora, CA, USA). Panel reactive antibody (PRA) and Single Antigen Anti-HLA antibodies were measured by the UCLA Immunogenetics Center (Los Angeles, CA, USA). Calculated PRA (cPRA) was estimated using the U.S. Organ Procurement and Transplantation Network (OPTN)'s online cPRA Calculator (https://optn.transplant.hrsa.gov/data/allocation-calculators/cpra-calculator/) and represents the percentage of organ donors that express 1 or more HLA antigens that transplant candidate had detectable anti-HLA antibodies against.

Methods for the detection of islet autoantibodies in subjects under the DEW-IT study have been previously described (*49*). Whole GAD65 or the intracellular portion of IA-2 were biosynthetically radiolabeled in the presence of $^{35}$S-methionine, while HPLC-purified $^{125}$I-monoiodo-Tyr$_{414}$-human insulin was purchased commercially from Perkin-Elmer (Hopkinton, MA, USA). Each antigen was incubated separately in triplicate wells with 2 to 5 uL of patient serum, then captured by Protein A, washed extensively to remove unbound antigen, and counted. Results were expressed using an index relative to specific positive control and negative control sera as described, with each cutoff defined as the 99$^{th}$ percentile of >200 individual healthy population controls. ZnT8 antibodies were detected via an analogous procedure using biosynthetically $^{35}$S-methionine radiolabeled antigen expressed from a construct comprising a hinged dimer containing the antigenic peptide loop of each of the aa325-Arg and the aa325-Trp alleles (JH6.2). Samples close to the cutoff, and those where triplicate well CV's were>30%, were reflexively repeated. The laboratory participates in DASP/IASP proficiency testing. The four antibody assays are each CLIA certified (CLIA# 50D0982418).

**Subhead 3: Plasma and platelet isolation from whole blood**

Platelets were isolated by centrifugation from blood, as previously described in (*50*). Briefly, whole-blood was centrifuged at 200 xg, 45° angle, for 15 minutes at room temperature to obtain platelet-rich-plasma (PRP). The PRP was transferred into Sarstedt Inc. Microvettes (Nümbrecht, Germany, cat# 20.1278.100) and centrifuged at 2550 xg for 5 minutes at room temperature. Plasma was aliquoted and stored at -80 °C until further use. Platelets were washed with DPBS and centrifuged at 2550 xg, subsequently lysed for SDS-PAGE and immunoblotting in 1% Nonidet P-40 (NP40) lysis buffer (25 mM HEPES pH 7.4, 1% NP40, 10% glycerol, 137 mM NaCl, 1 mM sodium vanadate, 50 mM sodium fluoride, 10 mM sodium pyrophosphate, 10 µg/ml aprotinin, 5 µg/ml leupeptin, 1 µg/ml pepstatin, and 1 mM phenylmethylsulfonyl fluoride). The platelet pellet was broken with a Becton Dickinson 1 ml 25g 5/8 syringe (Franklin Lakes, New Jersey, USA, cat# 309626) and centrifuged at max speed for 5 minutes at 4 °C and stored at -80 °C until further use.

**Subhead 4: Immunofluorescence staining**

Formalin-fixed paraffin-embedded (FFPE) pancreatic tissue sections from NOD versus NOR mice were prepared and assessed as previously described (*45*). To validate DOC2B antibody specificity for immunofluorescence imaging, pancreatic sections were immunostained with guinea pig anti-insulin [1:100; Thermo Fisher Scientific (Waltham, MA, USA), cat# A0564] and custom-made DOC2B antibody #2 [1:100; Abgent Inc. (San Diego, CA, USA)] extensively validated in (*32*), pre-incubated with blocking peptide (BP) constituting amino acids 96-116, the epitope of DOC2B antibody #2, custom made by Abgent Inc. (San Diego, CA, USA). The DOC2B antibody #2 (1:100) was mixed with BP (100x excess compared to antibody molarity) and rotated for 1 hour at 4°C before applying to the membrane to allow the blocking peptide to bind to the antibody. Alexa Fluor 647 goat anti-rabbit IgG (H+L) cross-absorbed secondary antibody (1:200; Thermo Fisher Scientific, cat# A21244) and 488 goat anti-guinea pig (H+L) secondary antibody (1:200; Thermo Fisher Scientific, cat# A11073) were used to detect DOC2B, and insulin, respectively. Slides were counterstained to mark the nuclei using 4',6-diamidin-2-phenylindole [DAPI; Vectashield Vector Laboratories (Burlingame, CA, USA)] and viewed using an LSM 900 confocal microscope (Carl Zeiss, Oberkochen, Germany). Confocal images were captured as a Z-stack using consistent acquisition parameters across all sections and then processed to generate a maximum intensity projection. Defined regions of interest (ROIs) were used to delimit the islets from acinar tissue.

**Subhead 5: Immunohistochemistry and insulitis scoring**

For the histological scoring of insulitis, slides containing formalin-fixed paraffin-embedded (FFPE) embedded whole pancreas were pre-treated with 3% peroxide [MilliporeSigma (Burlington, MA, USA), CAS# 7722-84-1] to remove endogenously expressed peroxidase from reacting with 3, 3'-diaminobenzidine (DAB) substrate kit from Vector Laboratories (Newark, CA, USA, cat# SK-4100). Subsequently, these were stained with guinea pig anti-insulin primary antibody, 1:50 dilution, from Invitrogen (Carlsbad, CA, USA, cat# PA1-26938) overnight at 4 °C; secondary goat anti-guinea pig IgG (H+L), biotin at 1:500 dilution, from Jackson ImmunoResearch Labs (West Grove, PA, USA, cat# 706-065-148) in blocking buffer; 5% Donkey serum (Jackson ImmunoResearch Labs, cat# 017-000-121), 0.1% Triton X-100 (MilliporeSigma, cat# X100-100ML) in 1X PBS, for 1 hour at room temperature. Samples were then incubated with Streptavidin peroxidase (Vector Laboratories, cat# SA-5004) at 1:250 dilution. DAB was added last to develop reaction. Samples were counter stained with Hematoxylin, Mayer's reagent from

Abcam (Waltham, MA, USA, cat#ab245880). Images were captured using a Keyence system (BZ-X710).

The score given for each mouse was based on the evaluation of all islets identified by morphology using a standard scale (*51, 52*) (0= no infiltrate, 1= non-invasive peri-infiltrate only, 2= infiltrating cells extending peri-islets <50%, 3= invasive insulitis >50%) as previously described (*53-55*).

**Subhead 1: Supplementary figures**

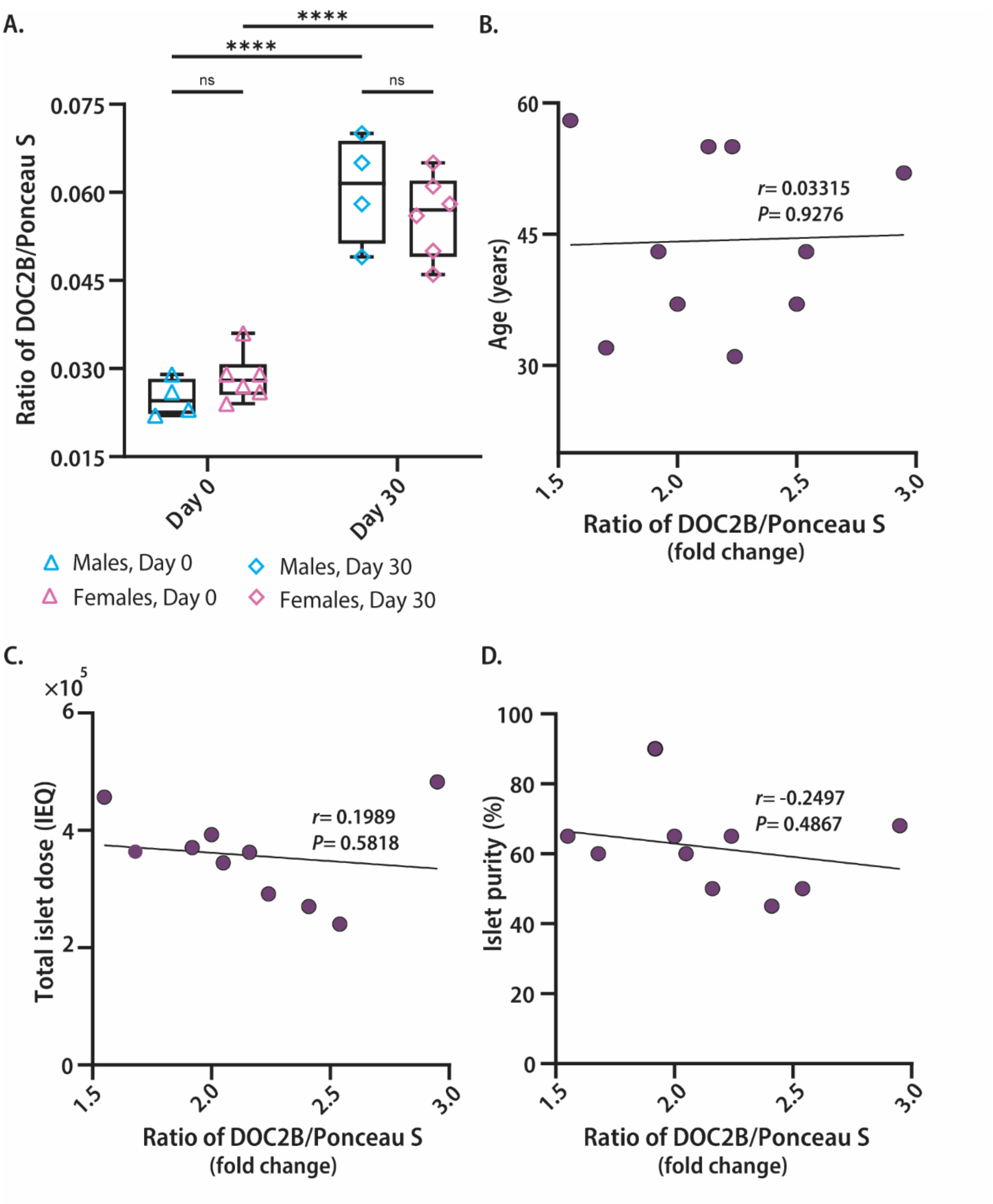


**Fig. S1. Plasma double C2-like domain beta protein levels by gender, and their correlation with age, total islet dose and islet purity in patients with long-standing T1D undergoing clinical islet transplantation.** (A) Plasma DOC2B levels (normalized to

Ponceau S) on Day 0 pre-, Day 30 post-transplantation were grouped by gender, n=*10* patients. Data are mean ± SEM. ****$P$<0.0001 by ordinary two-way ANOVA, using Tukey's post-hoc test. The fold change between plasma DOC2B levels (normalized to Ponceau S) on Day 30 post- versus Day 0 pre-transplant were correlated with (B) age of transplant patients, (C) total islets infused, or (D) islet purity, and are shown from $n$=10 individuals. There was no significant correlation between the fold change DOC2B levels on Day 30 post- versus pre-transplant (Day 0) and age [Pearson $r$= 0.03315, $P$= 0.9576]; total islet dose [Pearson $r$= 0.1989, $P$= 0.5818]; islet purity [Pearson $r$= -0.2497, $P$= 0.4867].

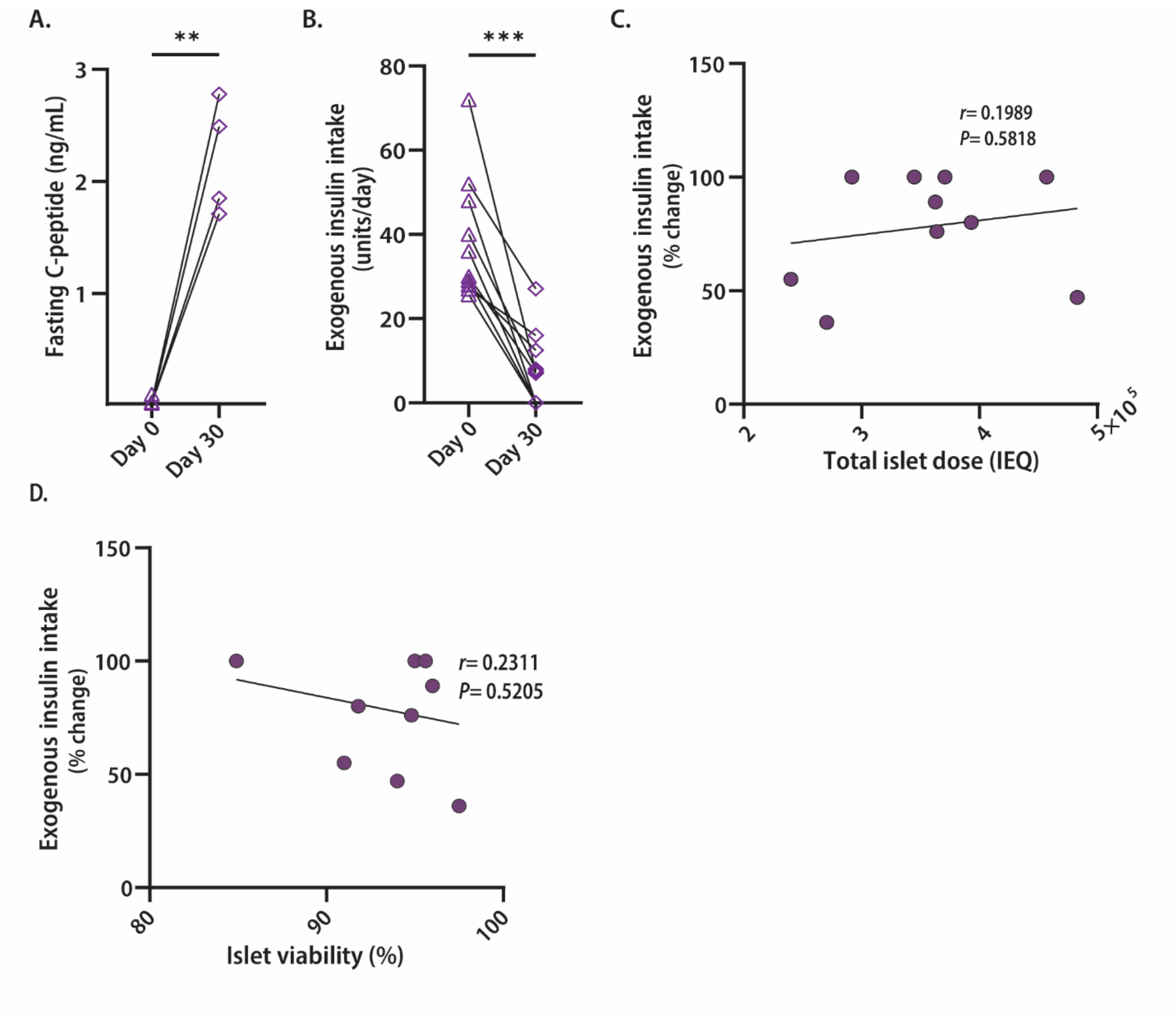


**Fig. S2. Clinical outcomes and their correlation with islet graft characteristics and plasma double c2-like domain beta protein levels in long-standing T1D patients pre- and post-islet transplantation**. Plasma from patients with long-standing T1D on Day 0 pre- and Day 30 post-transplantation was assessed for (A) fasting C-peptide levels in 4 individuals sampled pre-(Day 0) and post-(Day 30) islet transplant. Data represent the mean ± SEM; ***P*<0.01 using a two-tailed, unpaired, Welch's *t*-test. (B) Units per day of exogenous insulin intake on Day 0 pre- and Day 30 post-transplantation were assessed in 10 patients. Data represent the mean ± SEM; ****P*<0.001 using a two-tailed, paired,

Student's *t*-test. The percentage change of exogenous insulin intake on Day 30 post- versus Day 0 pre-transplant were correlated to (C) islet dose, (D) islet viability are shown. There was no significant correlation between the percentage change of exogenous insulin intake with islet dose or viability; insulin intake Day 30 post- versus pre-transplant (Day 0) and total islet dose [Pearson $r$= 0.1989, $P$= 0.5818]; islet viability [Pearson $r$= 0.2311, $P$= 0.5205].

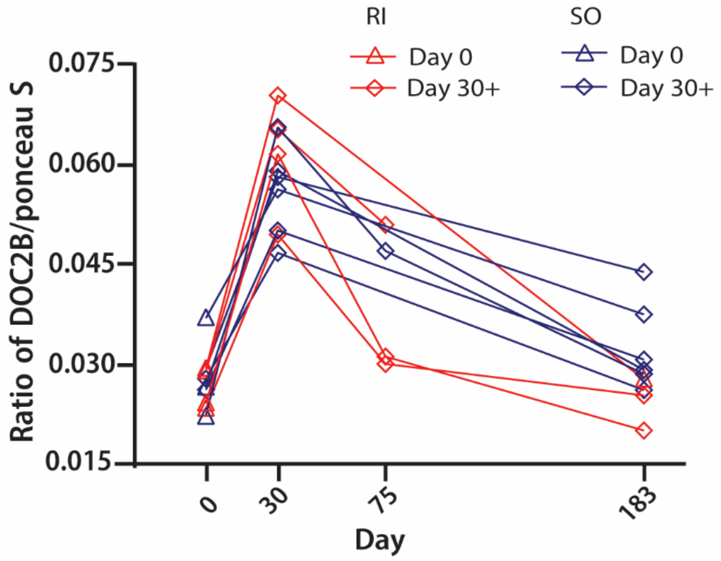


**Fig. S3. Longitudinal assessment of double C2-like domain beta protein levels in plasma from long-standing T1D patients pre- and post-islet transplantation**. Densitometry analysis of the plasma DOC2B levels normalization to Ponceau S (37-150 kDa range) from ten long-standing T1D patients pre-transplantation (Day 0, open triangles), and post-transplantation (Days 30, 75, 183; open diamonds). Patients were further grouped by those who resumed exogenous insulin intake (RI) on Day 75 and/or Day 183 (red), versus those who sustained an exogenous insulin independence outcome (SO) post-transplantation (blue). COH-027 RI on Day 75 and Day 183; COH-036 RI on Day 75, and COH-033 and COH-035 RI on Day 183.

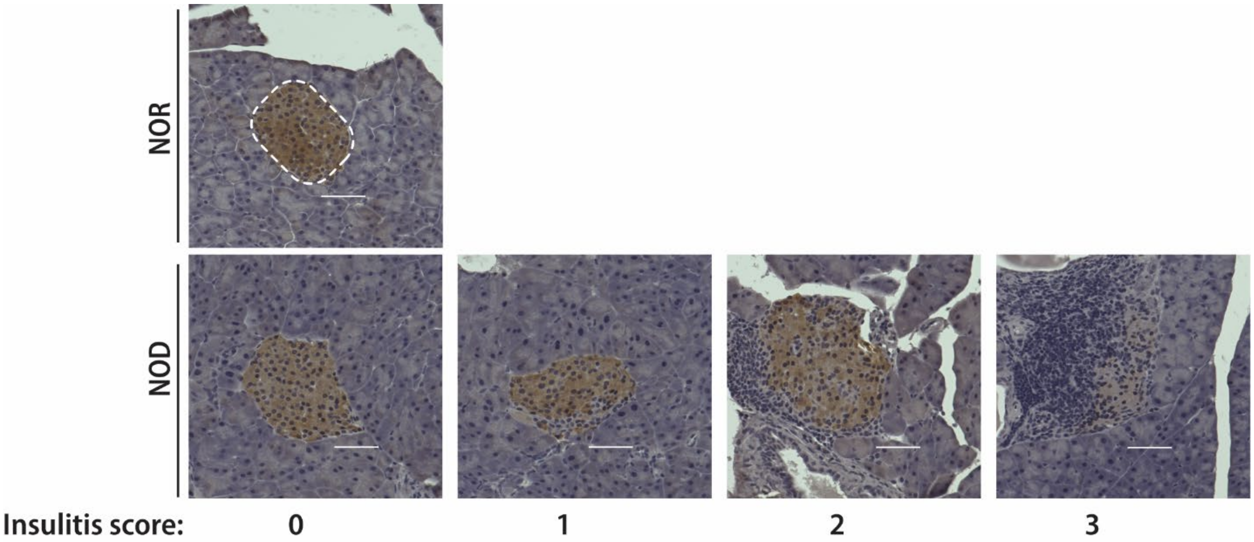


**Fig. S4. Insulitis scoring in 7-week-old female NOD and NOR mice pancreata.**

Representative images from n=4 independent measurements of insulitis scoring from hematoxylin-insulin-counterstained pancreatic tissue sections of 7-week-old female NOD and NOR mice. NOR control: enclosed within the white dashed line is a representative insulin-positive (brown) islet. Infiltrating leukocytes (dark blue). Scale bar= 20 µm.

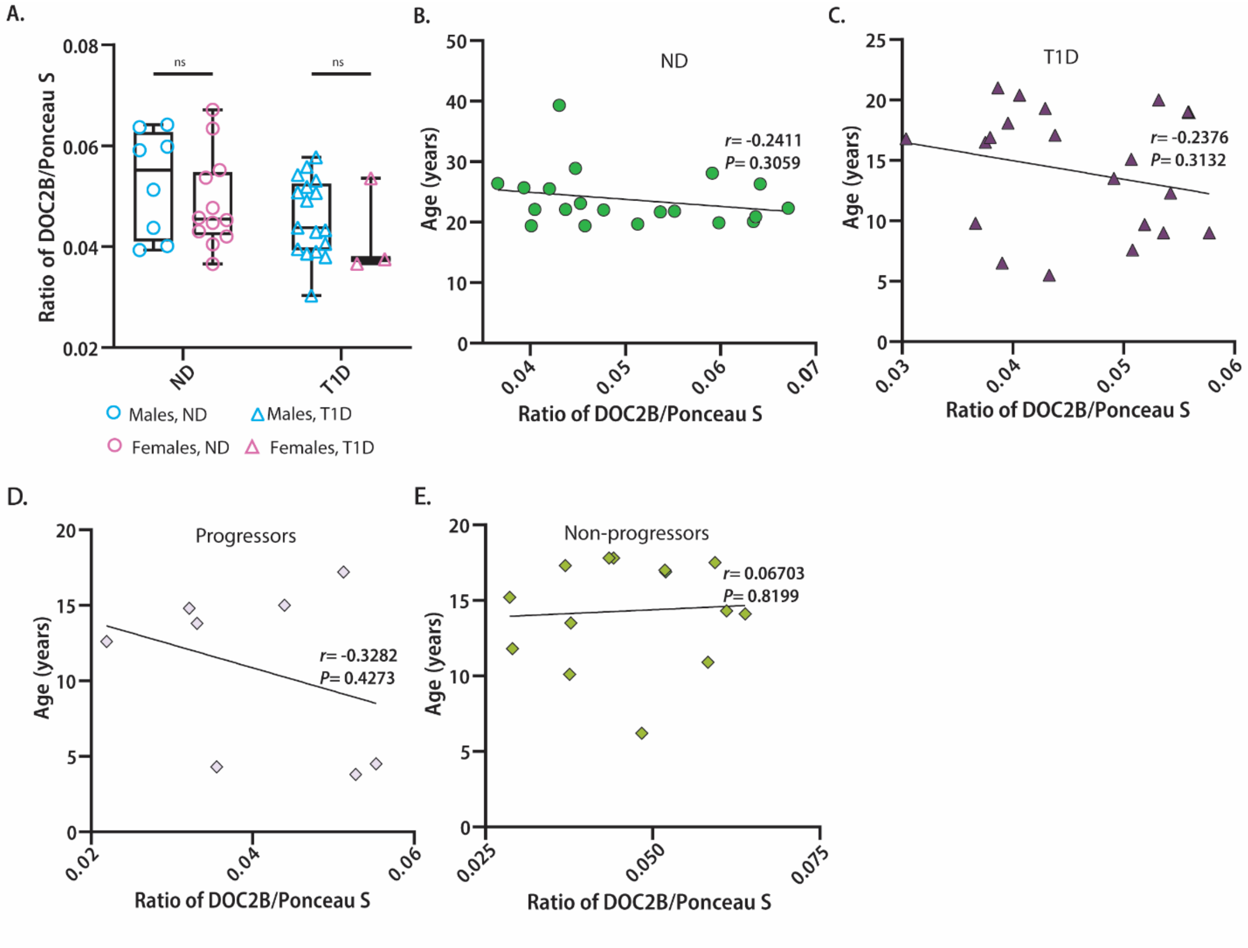


**Fig. S5. Double C2-like domain beta protein levels in males versus females, and their overall correlation to age in autoantibody-positive progressors to T1D or non-progressors, control T1D patients and ND individuals**. Plasma of autoantibody-positive individuals prior to diagnosis of their progression to T1D (progressors) or those that remained non-progressors, control T1D and individuals without diabetes (ND) from the DEW-IT study was assessed for DOC2B levels via quantitative immunoblot. (A) Densitometry analysis of the plasma DOC2B levels (normalized to Ponceau S) were grouped into males (light blue) and females (light pink) for ND controls (open circles) and individuals with T1D (open triangle), $n$=54 indviduals. Data represent the mean ± SEM. No significance (ns) by ordinary two way ANOVA, using Tukey's post-hoc test.

Age (B, C, D, E) in controls ND and T1D individuals, progressors or non-progressors to T1D were correlated with DOC2B levels (normalized to Ponceau S). There was no significant correlation between age and DOC2B levels across all groups evaluated. Age versus DOC2B levels in (B) control ND [Pearson $r$= -0.2411, $P$= 0.3059]; (C) T1D individuals [Pearson $r$= -0.2376, $P$= 0.3132]; (D) progressors [Pearson $r$= -0.3282, $P$= 0.4273]; (E) non-progressors [Pearson $r$= 0.06703, $P$= 0.8199].

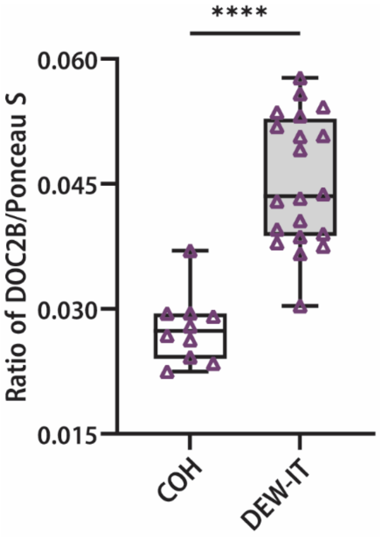


**Fig. S6. Plasma double C2-like domain beta protein levels are significantly lower in adult patients with long-standing T1D (COH) versus those in the pediatric DEW-IT cohort.** Densitometry analysis of plasma DOC2B levels normalized to Ponceau S (37-150 kDa range) from ten adult long-standing City of Hope (COH) T1D patients (10-52 year disease duration) sampled pre-transplantation (Day 0), versus T1D individuals (0-12.6 year disease duration) from the DEW-IT study, pediatric at enrollment. Individuals were not matched for comparisons. Data represent the mean ± SEM; n=10 subjects for the COH cohort and n=20 for the DEW-IT cohort; ****P<0.0001 using a two-tailed, unpaired, Welch's test.

### Subhead 3: References


49. V. Anand, Y. Li, B. Liu, M. Ghalwash, E. Koski, K. Ng, J. L. Dunne, J. Jönsson, C. Winkler, M. Knip, J. Toppari, J. Ilonen, M. B. Killian, B. I. Frohnert, M. Lundgren, A.-G. Ziegler, W. Hagopian, R. Veijola, M. Rewers, T. D. I. S. G. for the, Islet Autoimmunity and HLA Markers of Presymptomatic and Clinical Type 1 Diabetes: Joint Analyses of Prospective Cohort Studies in Finland, Germany, Sweden, and the U.S. *Diabetes Care.* **44**, 2269-2276 (2021).
50. A. Q. Reuwer, R. Nieuwland, I. Fernandez, V. Goffin, C. M. van Tiel, M. C. Schaap, R. J. Berckmans, J. J. Kastelein, M. T. Twickler, Prolactin does not affect human platelet aggregation or secretion. *Thromb. Haemost.* **101**, 1119-1127 (2009).
51. M. McDuffie, N. A. Maybee, S. R. Keller, B. K. Stevens, J. C. Garmey, M. A. Morris, E. Kropf, C. Rival, K. Ma, J. D. Carter, S. A. Tersey, C. S. Nunemaker, J. L. Nadler, Nonobese diabetic (NOD) mice congenic for a targeted deletion of 12/15-lipoxygenase are protected from autoimmune diabetes. *Diabetes.* **57**, 199-208 (2008).
52. G. Simon, M. Parker, V. Ramiya, C. Wasserfall, Y. Huang, D. Bresson, R. F. Schwartz, M. Campbell-Thompson, L. Tenace, T. Brusko, S. Xue, A. Scaria, M. Lukason, S. Eisenbeis, J. Williams, M. Clare-Salzler, D. Schatz, B. Kaplan, M. Von Herrath, K. Womer, M. A. Atkinson, Murine antithymocyte globulin therapy alters disease progression in NOD mice by a time-dependent induction of immunoregulation. *Diabetes.* **57**, 405-414 (2008).
53. M. J. Parker, S. Xue, J. J. Alexander, C. H. Wasserfall, M. L. Campbell-Thompson, M. Battaglia, S. Gregori, C. E. Mathews, S. Song, M. Troutt, S. Eisenbeis, J. Williams, D. A. Schatz, M. J. Haller, M. A. Atkinson, Immune depletion with cellular mobilization imparts immunoregulation and reverses autoimmune diabetes in nonobese diabetic mice. *Diabetes.* **58**, 2277-2284 (2009).
54. E. Oh, E. M. McCown, M. Ahn, P. A. Garcia, S. Branciamore, S. Tang, D. F. Zeng, B. O. Roep, D. C. Thurmond, Syntaxin 4 Enrichment in β-Cells Prevents Conversion to Autoimmune Diabetes in Non-Obese Diabetic (NOD) Mice. *Diabetes.* **70**, 2837-2849 (2021).
55. K. M. Bendtsen, C. H. Hansen, L. Krych, K. Buschard, H. Farlov, A. K. Hansen, Effect of Early-life Gut Mucosal Compromise on Disease Progression in NOD Mice. *Comp. Med.* **67**, 388-399 (2017).
